# Electrophysiological correlates of visual memory search

**DOI:** 10.1101/2024.05.17.594466

**Authors:** Lauren H. Williams, Iris Wiegand, Mark Lavelle, Jeremy M. Wolfe, Keisuke Fukuda, Marius V. Peelen, Trafton Drew

## Abstract

In everyday life, we frequently engage in ‘hybrid’ search, where we look for multiple items stored in memory (e.g., a mental shopping list) in our visual environment. Across three experiments, we used event-related potentials to better understand the contributions of visual working memory (VWM) and long-term memory (LTM) during the memory search component of hybrid search. Experiments 1 and 2 demonstrated that the FN400 – an index of LTM recognition – and the CDA – an index of VWM – increased with memory set size (target load), suggesting that both VWM and LTM are involved in memory search, even when memory load exceeds capacity limitations of VWM. In Experiment 3, we used these electrophysiological indices to test how categorical similarity of targets and distractors affects memory search. The CDA and FN400 were modulated by memory set size only if items resembled targets. This suggests that dissimilar distractor items can be rejected before eliciting a memory search. Together, our findings demonstrate the interplay of VWM and LTM processes during memory search for multiple targets.

## Memory in multiple-target search

When shopping at the grocery store, you likely have a shopping list of multiple items (e.g., noodles, spaghetti sauce, and bread) you intend to purchase. This routine task involves a “hybrid” search through memory (i.e., your mental shopping list) and through the visual environment (i.e., the items at the grocery store)(Wolfe, 2012, 2021). Hybrid search is also an important task for many professionals, such as radiologists and baggage screeners, who need to keep an eye out for a wide-range of possible abnormalities or hazardous items in an x-ray (Wolfe, Alaoui Soce, & Schill, 2017). Although hybrid search is ubiquitous in daily life and observers can search for as many as 100 unique items quite effectively (Wolfe, 2012), relatively little is known about the neural mechanisms that support memory search during search for multiple targets (Ort & Olivers, 2020).

In hybrid search tasks, response time increases linearly *with the log* of the number of search targets stored in memory (memory set size, MSS), suggesting that target verification is highly efficient (Wolfe, 2012). During search for multiple targets, one could imagine that multiple “search templates” are pre-activated in visual working memory (VWM), biasing attention towards target-relevant features in the environment (Desimone & Duncan, 1995). Then, at a post-selection stage, the observer determines whether the attended item matches any of their pre-activated search templates (Ort & Olivers, 2020). This use of VWM would be limited by the capacity of that VWM, which is assumed to be around 4 items (Luck & Vogel, 1997). However, when the number of targets exceeds the small capacity limitations of VWM, target verification was suggested to rely mostly on long-term memory (LTM) recognition (Ort & Olivers, 2020). LTM appears to be essentially free of capacity limitations for targets that are sufficiently learned (Brady et al., 2008). Support for this comes from hybrid search studies, in which observers memorized up to 100 targets. Eye-tracking data showed that observers searched the entire display and performed a memory search for each selected item until the actual target was found (Drew, Boettcher, & Wolfe, 2017). Furthermore, performance was not impaired if a concurrent task was used to “fill up” VWM capacity during search (Drew, Boettcher, & Wolfe, 2016). Though search was not impaired, VWM capacity seemed to be reduced by roughly one item when performing a hybrid search task concurrently with a VWM load. This evidence suggests that the target load in hybrid search does not fill up VWM. Rather, VWM may serve as a fixed-capacity conduit that passes a single attended item at a time into long-term memory for target verification.

While it seems clear that the multiple templates used in hybrid search do not occupy VWM, other recent work, provides strong evidence that VWM might contain templates that can guide hybrid search even at large MSSs (Cunningham & Wolfe, 2014). This would occur, not by having dozens of search templates in VWM, but by having a few “guiding templates” in VWM that would make item selection more efficient while dozens of “target templates” in LTM would be used to determine if a selected item was indeed a target (Wolfe, 2021). For example, imagine a hybrid search task in which your targets are from the same category (e.g. all targets are animals; perhaps, a cat, a cow, and an octopus), a rough representation of the features of animals could guide attention *away* from items that share few visual features with the target category (e.g., flags). These items could be rejected before eliciting memory search. Other items, other animals or distractors with enough similar features could be attentionally selected and would require memory search to determine if they were a member of the memory set (Cunningham & Wolfe, 2014). A recent study provided behavioral and neural evidence for such top-down attentional effects of categorical target templates during hybrid search (Shang et al., 2024). Specifically, distractors from the same category elicited a stronger N1 and N2 posterior-contralateral (N2pc) than distractors from a different category. The N2pc is a lateralized posterior event-related potential (ERP) occurring around 200 ms after the stimulus’ onset, marking spatial attentional selection (Eimer, 1996; Luck & Hillyard, 1994b; Wiegand et al., 2018). While these results highlight the role of feature-based attentional selection in multiple-target search, it remains unclear how the number of targets, and their categorical level, influences VWM and LTM processes following attentional selection.

## Electrophysiological correlates of memory processes during multiple-target search

Further insights about the memory processes in hybrid search may be derived from ERPs marking post-selective processing. ERP correlates of LTM for large sets of learned items have been studied extensively in the recognition memory literature. Here, previously learned “old” probe items are known to elicit two dissociable ERP components, compared to new items. First, the FN400 occurs 300-500 ms after probe. It is fronto-centrally distributed and less negative (i.e. more positive) for old items than for new items. The FN400 is thought to reflect familiarity-based recognition, a fast-acting process that does not reflect retrieval of qualitative or contextual details about the encoding event (Curran, 2000). Second, the Late Positive Complex (LPC) occurs around 500 ms after the probe. It is centro-parietally distributed, and more positive for old than for new items. The LPC is thought to reflect recollection, the more effortful retrieval process for the recognized item along with its context of occurrence (Rugg & Curran, 2007). The FN400 and LPC are elicited for different memoranda, including visual objects (Curran & Cleary, 2003). Thus, the FN400 and/or LPC may also serve as markers of LTM retrieval during target verification in hybrid search.

The Contralateral Delay Activity (CDA) is a posterior, lateralized sustained negative ERP that occurs during VWM maintenance ∼300-1000 ms after the presentation of to-be-remembered items (Luria et al., 2016). In change-detection VWM tasks, the CDA increases with the number and resolution of maintained items, reaching an asymptote at participants’ individual VWM capacity limit (Fukuda, Awh, & Vogel, 2010; Ikkai, McCollough, & Vogel, 2010; Vogel & Machizawa, 2004; Wiegand et al., 2014). Hence, the CDA is regarded as a neural signature of active storage in VWM. In visual search, a CDA (also called sustained posterior contralateral lateralization, SPCN) follows the N2pc if the task requires VWM maintenance for post-selective target discrimination (Brisson & Jolicoeur, 2008; Hilimire et al., 2011; Wiegand et al., 2013). The CDA has also been used as a neural signature of pre-activating attentional templates in VWM using simple stimuli and small set sizes of one to three targets (Carlisle et al., 2011; Grubert, Carlisle, & Eimer, 2016; Gunseli, Meeter, & Olivers, 2014; Luria & Vogel, 2011). However, whether the CDA also marks capacity-limits within VWM during memory search for multiple targets has not been investigated, yet.

## The present study

The goal of the present study was to better understand the contributions of different memory processes during the post-selection stage of multiple-target search (Ort & Olivers, 2020). In three experiments, we measured ERP correlates of VWM (CDA), and LTM recognition (FN400 and LPC) in a modified hybrid search task, where visual set size was constrained to only one task-relevant item per trial, to isolate the effects of searching through memory rather than searching through space. The processes contributing to memory search, and their hypothesized ERP correlates, are illustrated in Figure 1. Across all experiments, we varied the MSS, that is, the number of targets observers would search for. One of the targets was present in half of the trials and a distractor was displayed in the other half. In Experiments 1 and 2, observers searched for sets of 1-64 distinct, realistic objects. We tested whether and how the CDA, FN400, and LPC would vary with MSSs, marking the contributions of VWM and LTM processes for target load within and beyond the capacity limitations of VWM. In Experiment 3, observers searched for 2 or 16 target objects either among distractors of the same, a similar, or dissimilar category. We tested whether target dissimilar items would indeed be rejected before eliciting a memory search (Cunningham & Wolfe, 2014), as marked by the ERP correlates of memory search established in Experiments 1 and 2.

**Figure 1:**
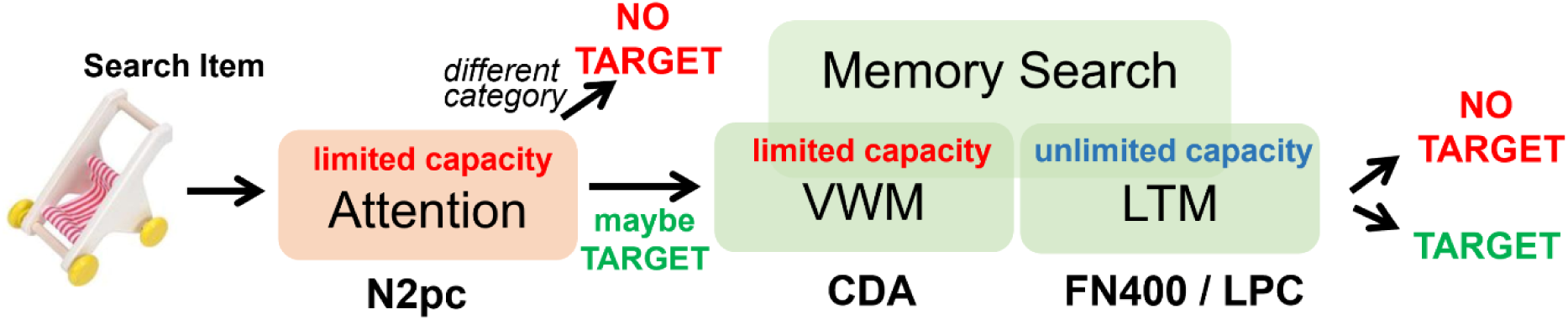
Schematic illustration of processes involved in memory search and their hypothesized ERP correlates. We assumed that the CDA, indexing visual working memory (VWM) load (Vogel & Machizawa, 2004), would increase with memory set size (MSS) up to the VWM capacity-limit of ∼4 items. By contrast, we hypothesized that the FN400 and/or LPC, marking the strength of long-term memory (LTM)-based recognition (Rugg & Curran, 2007), would vary with MSS beyond 4 items. Finally, we expected that no memory search would be elicited if search items can be rejected on an attentional stage based on categorical features (Shang et al., 2024); thus, items dissimilar to any of the targets would not be loaded into VWM or LTM.

## Experiment 1

In Experiment 1, observers searched for either 1, 2, 4, or 8 distinct target objects to distinguish between MSS effects on ERPs within and beyond the capacity limitations of VWM. We analyzed response times and accuracy, together with the N2pc, CDA, FN400 and LPC as a function of MSS and target presence. If a limited number of pre-activated search templates within capacity limitations of VWM supports target verification (Ort & Olivers, 2020), we may expect the CDA to increase with MSS but plateau at MSS4. Alternatively, if VWM indeed only passes the attended item into LTM for target verification (Drew et al., 2016, 2017), a CDA should be present, but should not be sensitive to MSS. For target sets exceeding the small capacity limitations of VWM, we expected that target verification would rely largely on LTM recognition (Ort & Olivers, 2020), which would be reflected in a modulation of the FN400 and LPC by target presence (i.e. old/new difference) and MSS beyond 4 items. Finally, we expected that an N2pc should be elicited, marking spatial attentional processing of target items. Note that spatial attention to the target location was already pre-cued by color in the present design; thus, the N2pc cannot be interpreted to reflect template-based guidance to target features in the presence of distractors. Rather, the N2pc may reflect attentional processing of an item *at* the attended location, which is expected to be easier if one, or a few, search templates can be activated compared to when many targets are in the memory set.

### Method

#### Participants

Twenty-six participants from the University of Utah participated in Experiment 1 for course credit or $15.00 an hour. Four participants had artifact rejection-rates that exceeded our exclusion criteria (30%) and the full EEG session was inadvertently not recorded for two participants, leaving a total of 20 participants in the final dataset (15 female / 5 male, average age: 22.4 years, age range: 18 to 34 years). A sample size of 20 was pre-registered (https://osf.io/saz82). The study was approved by the University of Utah’s Institutional Review Board, and all participants provided informed written consent.

#### Memory Search Task

Participants completed a memory search task for sets of 1, 2, 4, or 8 (MSS) unique target items (Figure 2), which were sampled randomly for each participant from Brady et al. (2008). For each MSS, participants completed a task block with three phases: memorization, recognition test, and memory search. During the memorization phase, participants memorized the items in their target list as they were displayed one at a time at the center of the screen for 3000 ms. Next, participants completed an old/new recognition memory test to ensure the targets had been memorized. The target objects and an equal number of foils were presented one at a time at the center of the screen, and participants indicated whether or not the object was a member of the target memory set. In order to proceed to the next phase of the experiment, participants were required to pass the recognition test with a score of 100% twice. Participants were allowed five attempts to complete the testing phase, and no participant failed to meet this criterion. Finally, participants completed the critical memory search phase. At the beginning of each search block, participants were instructed to attend to the item cued in either a blue or green frame (counterbalanced across participants). At the beginning of each trial, a blank screen containing only a fixation cross was displayed for 300 – 400 ms. To measure lateralized ERP components, the attended item was displayed to the left or right side of the fixation cross in the task-relevant color (e.g., blue), and an unattended item was shown on the opposite side of the fixation screen in the task-irrelevant color (e.g., green). The objects appeared for 200 ms, followed by a response interval where only the fixation cross was visible. Participants indicated whether or not the object was a member of the target set using the ‘F’ or ‘J’ keys. For each MSS, there were 400 trials (50% target present, 50% target absent). The response screen was displayed for either 1000 ms or until participants made a response. Finally, participants were shown feedback on their performance until they pressed the space bar to start the next trial. The order of blocks was counterbalanced.

**Figure 2.**
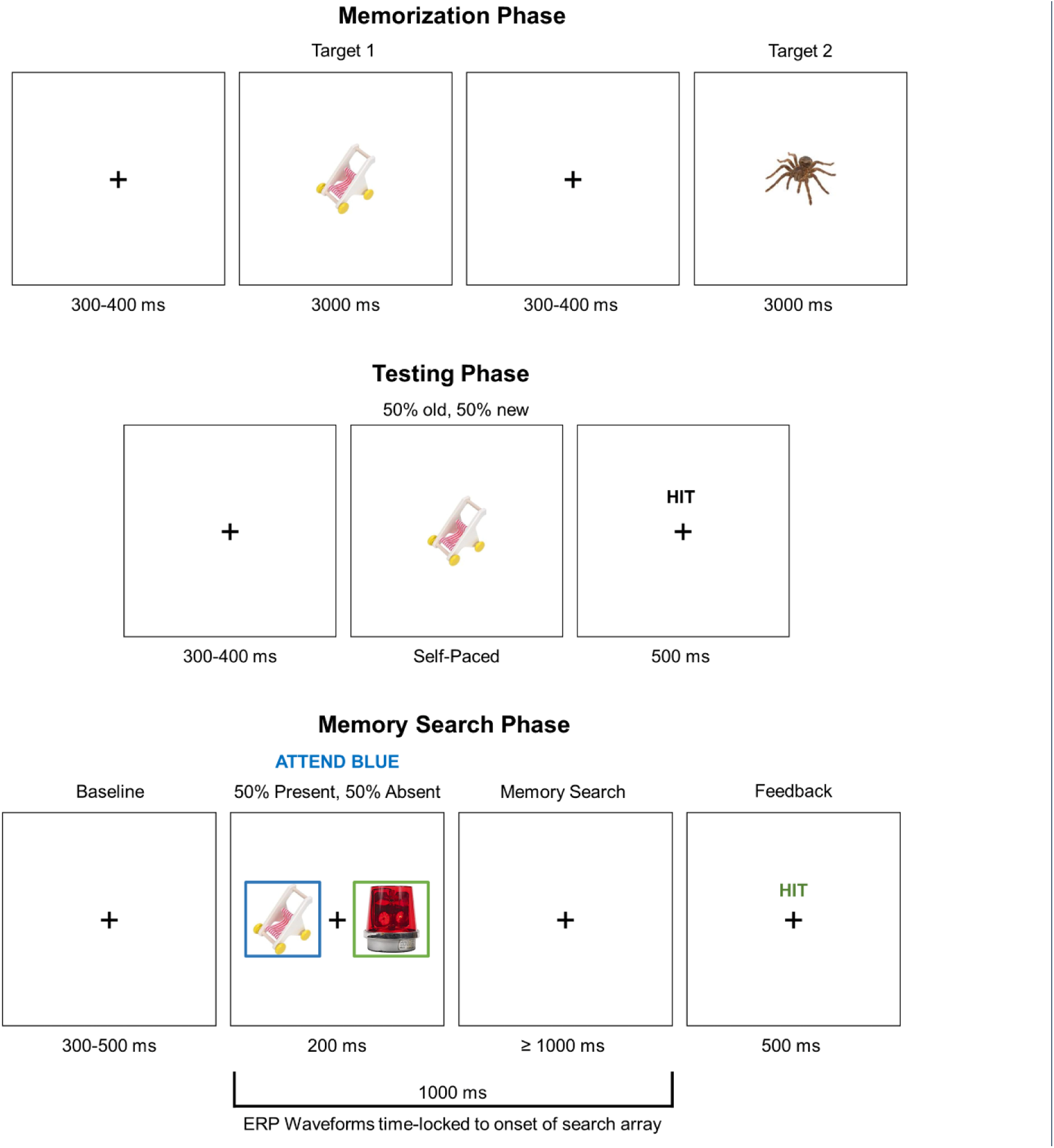
Experimental design for Experiments 1 (Memory Set Size, MSS 1, 2, 4, & 8) and 2 (MSS 2, 4, 16, & 64). The passing rate of the memory test phase was 100% in Experiment 1, and 80% or higher in Experiment 2.

#### EEG Procedure

EEG data was recorded at 500Hz with Brain Products’ ActiCap and ActiCHamp system using 32 active electrodes from the International 10/20 system. The data were referenced online to the average of the left and right mastoids. Electrode impedance was reduced to 15 kOhms or lower at each electrode site prior to recording Twenty-eight channels besides mastoids were placed according to the 10-20 system with 2 additional placed on the outer canthi bilaterally to recorder HEOG and Fpz serving as the ground. The channels included Fz, Cz, Pz, and Oz, as well as Fp1, F7, F3, FC5, FC1, T7, C3, CP5, CP1, P7, P3, PO7, and contralateral homologues.

#### EEG Analysis

The EEG data was processed offline using the EEGLAB (Delorme & Makeig, 2004) and ERPLAB (Lopez-Calderon & Luck, 2014) toolboxes for MATLAB. First, a high-pass filter at 0.01 Hz with a half-amplitude cutoff was applied to the raw EEG data. Next, the data was epoched from -200 – 1000 ms and trials with artifacts were removed from the analysis (see below). The stimulus-locked ERP waveforms were time-locked to the onset of the memory search and extended for 1000 ms. The response-locked waveforms (see Supplementary Analyses) were time-locked to the participant’s response and went backward in time for 1000 ms. Both the stimulus-locked and the response-locked waveforms were baseline-corrected to the 200 ms prior to the onset of the memory search. For eye-movement detection, two electrodes were placed ∼1 cm from the external canthi of each eye, and an HEOG channel was created offline by subtracting the left and right eye channels. A step-function was then applied to the HEOG channel in order to detect eye-movements (threshold 40 µV). Blinks and other large artifacts were detected using a moving-window function (threshold 140 µV) applied to the frontal electrodes (Fp1/2) above each eye. Individual thresholds were adjusted for each participant as needed in order to increase the signal to noise ratio. On average, 9.7% of trials were excluded for the stimulus-locked waveforms and 9.3% of trials were excluded for the response-locked waveforms. Finally, we applied a low-pass Butterworth filter with a half-amplitude cutoff of 30 Hz for plotting purposes only. The statistical analyses were performed on the data prior to the application of the low-pass filter.

#### Event-related potentials

The selection of electrodes and time windows was pre-registered (https://osf.io/saz82) and in accordance with previous studies of the N2pc (Drew et al., 2018; Eimer, 1996; Luck & Hillyard, 1994a; Vogel & Machizawa, 2004), and FN400 and LPC, specifically for using object images (Drew et al., 2018; Küper et al., 2012; Küper & Zimmer, 2018)

For the non-lateralized components, the FN400 and LPC, we used the average of electrodes Fz, Cz, Pz, F3/4, C3/4, and P3/4. The mean amplitude of the FN400 was measured between 300-500 ms relative to stimulus onset, and the mean amplitude of the LPC was measured from 500-800 ms relative to stimulus onset. In addition, we created an old-new difference wave by subtracting the target absent trials from the target present trials and measured the mean amplitude and 50% fractional area latency of the waveform in the time window 300-800 ms (i.e., the point at which the area reached 50% of the total area between 300–800 ms).

For the lateralized components, the N2pc and CDA, we created a contralateral-ipsilateral difference wave relative to the attended side of the display using the average of electrodes P07/08 and P7/8. The mean amplitude of the N2pc was measured between 200-300 ms. The mean amplitude of the CDA was measured between 300-1000 ms. We also analyzed the response-locked CDA (-300 to 0 ms) to ensure any observed differences in mean amplitude between conditions were not driven by differences in response-time (Ankaoua & Luria, 2023; Williams & Drew, 2021), which are reported in Supplementary Analyses. Overall, the results of the analyses on the response-locked CDA mirrored our findings in the stimulus-locked analyses; the response-locked

#### Statistical Analysis

The analyses for Experiment 1 were pre-registered (https://osf.io/saz82), except the analyses of the fractional area latency. The ERP and response-time analyses were performed for correct trials only. For each dependent measure, we conducted a 2 (present, absent) by 4 (MSSs 1, 2, 4, 8) repeated-measures ANOVA. This analysis differed from the pre-registered plan to evaluate MSS effects using separate one-way ANOVAs for target present and target absent trials to look for interactions between target presence/absence and MSS. None of the MSS effects substantively differed if we instead use the pre-registered analysis. In addition to frequentist statistics, we computed Bayes Factors for each analysis in order to quantify the degree of evidence for the alternative relative to the null hypothesis (BF_10_). Bayes Factors greater than 3 were considered sufficient evidence for the alternative hypothesis, and Bayes Factors less than 1/3 were considered sufficient evidence for the null hypothesis (Jeffreys, 1998). Effects and interactions were followed up with Tukey’s tests for multiple comparisons.

### Results

#### Response Time and Accuracy

Response time increased with MSS, F(3, 57)=7.66, p<.001, η^2^=.29, BF_10_=410531.8 (Figure 3a). Significant differences were found for set sizes 1 vs. 4, 1 vs. 8, and 2 vs. 8, all p<.01. Responses were faster for target present trials than target absent trials, F(1, 19)=32.18, p<.001, η^2^=.63, BF_10_=30.65. The MSS by target presence interaction was not statistically significant, F(3, 57)=0.38, p=.77, η^2^=.02, BF_10_=.075. Accuracy (percent correct) did not significantly differ with MSS, F(3, 57)=0.82, p=.49, η^2^=.04, BF_10_=.14 (Figure 3b). Target present trials were less accurate than target absent trials (i.e. more misses than false alarms), F(1, 19)=22.53, p<.001, η^2^=.54, BF_10_=137.25. The set size by target presence interaction was not statistically significant, F(3, 57)=2.57, p=.06, η^2=^.12 , BF_10_=.25.

**Figure 3.**
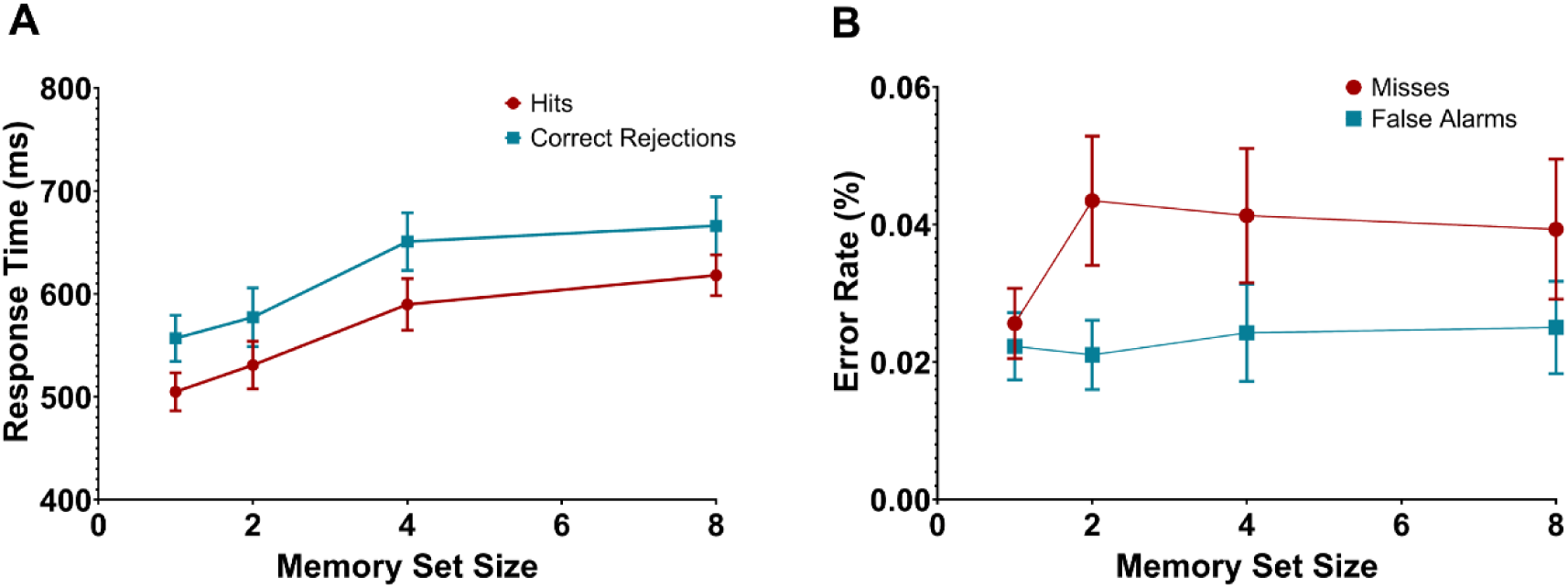
Behavioral data for Experiment 1: A) response time. B) error rate. Error bars represent the standard error of the mean throughout the manuscript.

#### FN400, LPC, and old-new effect

First, we evaluated the non-lateralized waveforms (Figure 4). If memory search relies on LTM recognition, we would expect that the FN400 and LPC would be more positive if observers recognize “old” targets compared to when being presented with a distractor item. Consistent with our prediction, both, the FN400 and the LPC were more positive in target present than target absent trials, F(1, 19)=95.9, p<.001, η^2^=.83, BF_10_=4.51e+20 and F(1, 19)=104.3, p<.001, η^2^=.85, BF_10_=1.55e+18.

**Figure 4.**
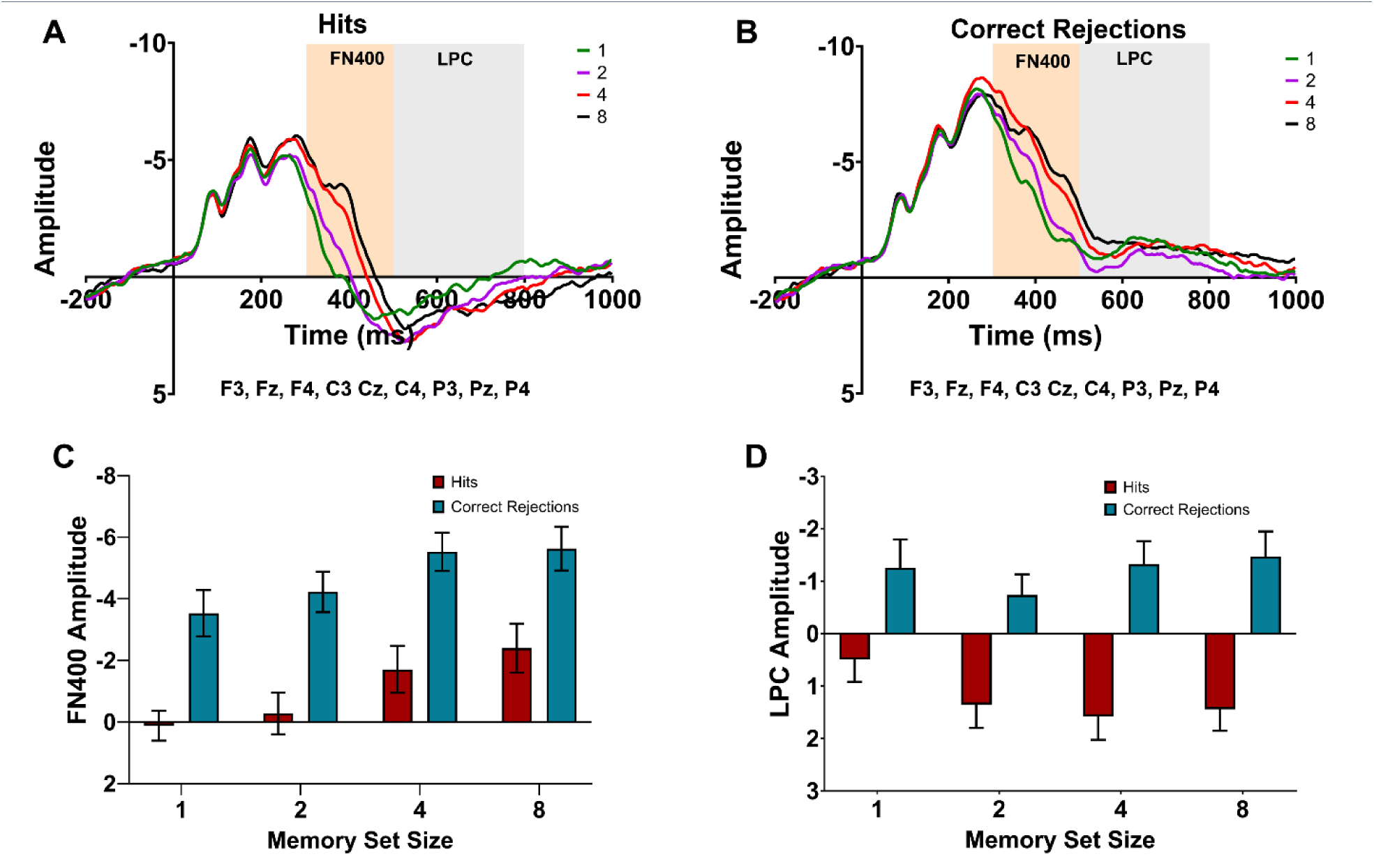
Non-lateralized data for Experiment 1: A) non-lateralized waveforms for target present trials. B) non-lateralized waveforms for target absent trials. C) mean FN400 amplitude. D) mean LPC amplitude.

Furthermore, we expected the recognition signal to be stronger when a smaller set of targets needed to be distinguished from distractors compared to a larger set. This was true for the FN400. The FN400 varied with MSS, F(3, 57)=16.22, p<.001, η^2^=.46 , BF_10_=192.23 (Figure 4c) and was significantly smaller (less negative) in amplitude for set size 1 than set size 4, p<.001, and set size 8, p<.001, and smaller for set size 2 than set size 4, p=.004, and set size 8, p<.001. There were no significant differences between any of the other MSSs, all p-values>.05. The set size by target presence interaction was not statistically significant, F(3, 57)=0.78, p=.51, η^2^=.04 , BF_10_=.10; thus, the FN400 was modulated by MSS across target present and absent trials (i.e. hits and correct rejections). Different from the FN400, the LPC did not differ between MSSs, F(3, 57)=1.27, p=.29, η^2^=.06 , BF_10_=.10 (Figure 4d) when target present and absent trials were combined. However, there was a significant set size by target presence interaction, F(3, 57)=5.34, p=.003, η^2^=.22 , BF_10_=.54. Only in target present trials, the LPC was less positive for set size 1 than for set size 2, p=.007, and for set size 4, p=.002). No significant differences in the LPC between set sizes were found in target absent trials (all p-values>.05).

Finally, the amplitude of the old-new difference waves (ERP in response to target-present trials minus ERP in response to target-absent trials across the time windows of the FN400 and LPC, Figure 5a), did not vary with MSS, F(3, 57)=1.60, p=.20, η^2^=.08, BF_10_=.37 (Figure 5b). However, the fractional area latency of the old-new difference waves significantly varied with MSS, F(3, 57)=4.93, p=.004, η^2^=.21, BF_10_=11.77 (Figure 5c). MSS 8 had a significantly longer fractional area latency than MSS 1, p=.02, and MSS 2, p=.006.

**Figure 5.**
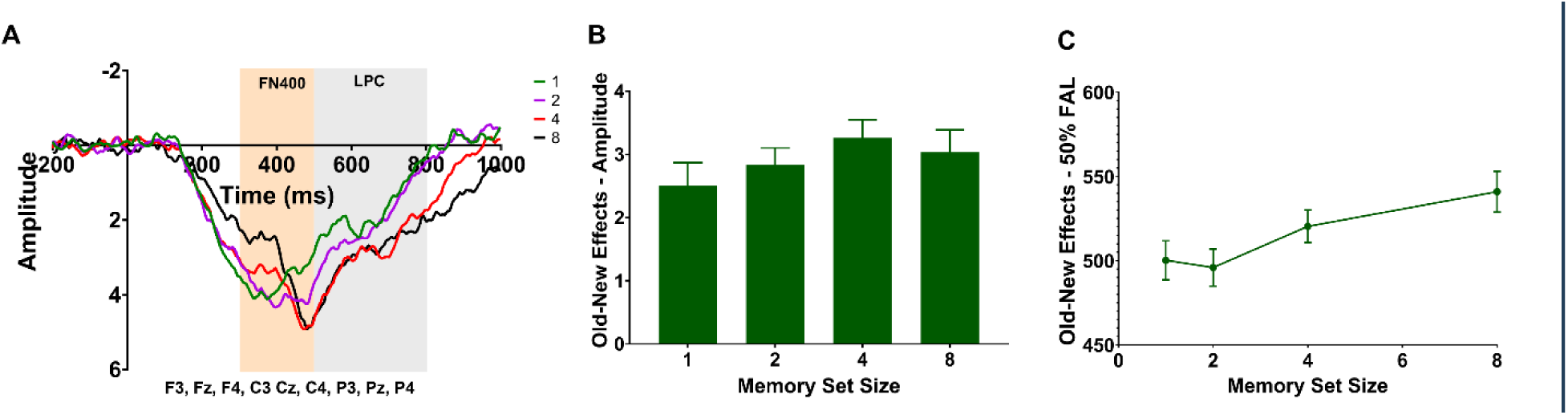
Old/new difference waves for Experiment 1: A) old/new difference waveforms B) mean amplitude of the old/new difference wave C) mean fractional area latency of the old/new difference wave

#### CDA

Next, we evaluated the stimulus-locked CDA (Figure 6). We expected that the CDA amplitude would increase with MSS up to the capacity limitation of VWM of ∼4 items, but not further. Indeed, we did find that the amplitude of the CDA varied with set size, F(3, 57)=4.15, p=.01, η^2^=.18, BF_10_=.57 (Figure 6c). Furthermore, target present trials had a larger CDA amplitude than target absent trials, F(1, 19)=9.8, p=.006, η^2^=.34, BF_10_=3518.45, and the MSS by target presence interaction was significant, F(3, 57)=6.16, p=.001, η^2^=.24, BF_10_=9.15, reflecting the fact that the CDA did not vary with MSS for target absent trials, p=.53. Interestingly, the target present CDA amplitude was larger at MSS 8 than MSS 4, p=.009. Thus, different from our prediction, the CDA increased beyond the usual capacity limitations of VWM, rather than showing a plateau after set size 4.

**Figure 6.**
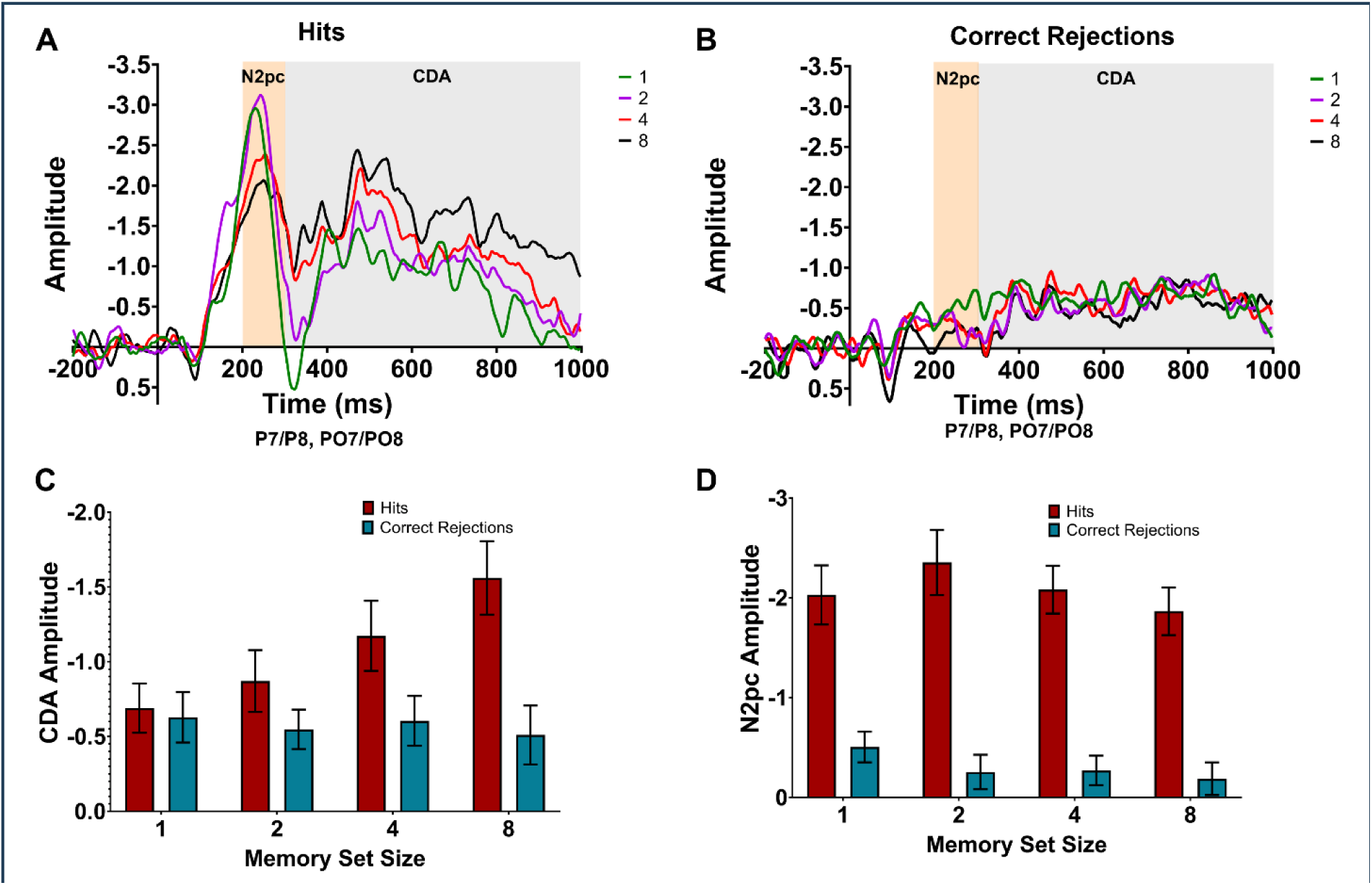
Lateralized stimulus-locked data for Experiment 1: A) contralateral-ipsilateral waveforms for target present trials. B) contralateral-ipsilateral waveforms for target absent trials. C) mean N2pc amplitude. D) mean CDA amplitude.

**Figure 7.**
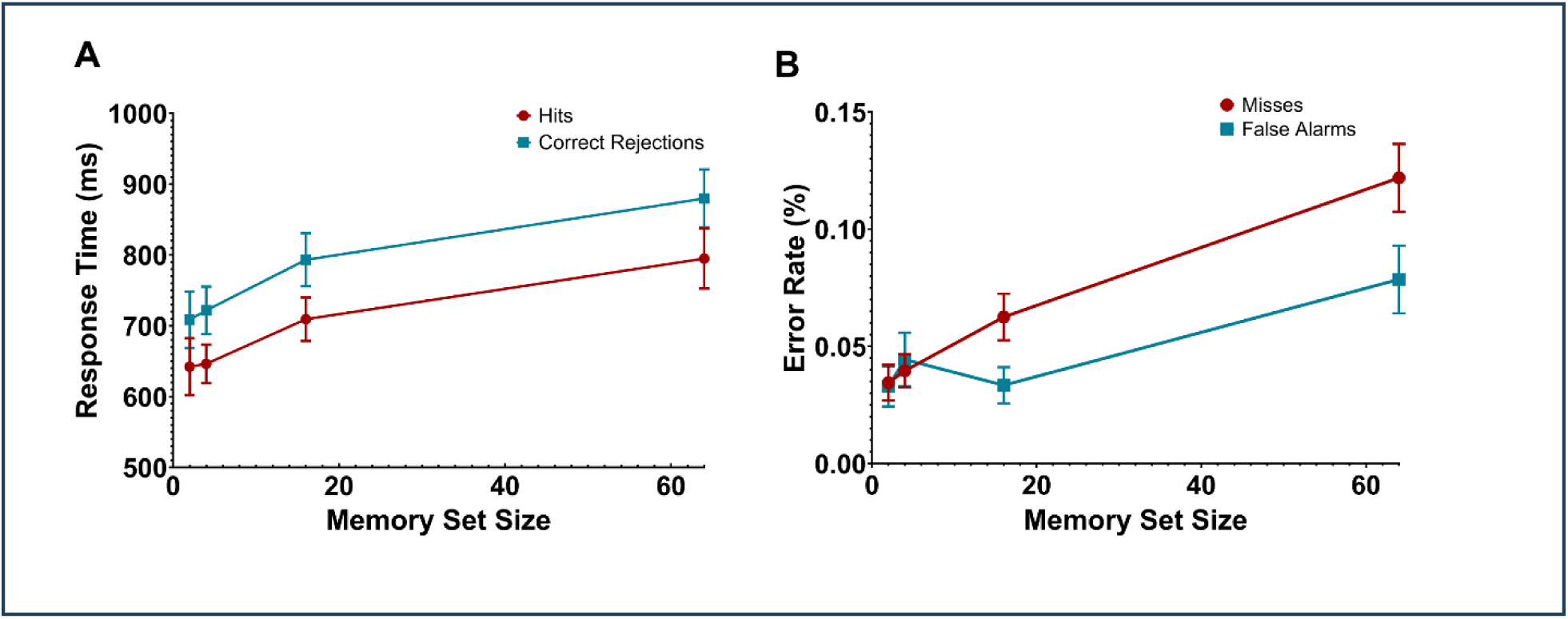
Behavioral data for Experiment 2: A) response time. B) error rate.

#### N2pc

Finally, we analyzed the N2pc, marking attentional processing of the selected item. The N2pc amplitude was larger for target present than target absent trials, F(1, 19)=113.6, p<.001, η^2^=.86, BF_10_=2.33e+27. While the visual inspection of the grand-averaged N2pc suggests that the amplitude was indeed higher for MSSs 1 and 2 compared to 4 and 8 in target present trials, the amplitude of the N2pc did not significantly vary with MSS, F(3, 57)=1.26, p=.30, η^2^=.06, BF_10_=.05 (Figure 4c). Neither was the set size by target presence interaction statistically significant, F(3, 57)=1.26, p=.30, η^2^=.06, BF_10_=.21.

### Discussion

In Experiment 1, we found that ERP correlates of both VWM and LTM recognition varied with target load beyond set sizes of four targets during memory search. First, the FN400 and LPC were more positive if a target was present, showing the expected old/new effect reflecting item recognition (Curran, 2000). Further, there was an effect of MSS on the amplitudes of the FN400 and the latency of the old/new effect. The FN400 is thought to reflect item familiarity, and has been related to perceptual matching processes when comparing visually processed information to memory representations (Küper et al., 2012; Zimmer & Ecker, 2010). The effect of target load suggests that the familiarity signal was weaker when more candidate items needed to be retrieved from LTM in order to be compared to the attended item in the search display during memory search. The prolonged latency of the old/new difference waves with larger MSS may reflect the increase in time required for matching more items of the memory set against the attended item. By contrast, the LPC, marking recollection of items and their study context (Norman et al., 2008), varied little between MSSs. This suggests that differences between MSS conditions do not result from recollection of the study episode. This is in line with the rather small response time costs of increasing the target load, supporting the assumption that memory search is a fast-acting matching process, which does not rely on time-consuming recollection of each item in the memory set (Nosofsky et al., 2014); however, see (Wiegand & Wolfe, 2020; Wolfe et al., 2015).

In a deviation from our predictions, the CDA significantly increased up to MSS 8. Previous research on hybrid search suggested that VWM usage does not increase with MSS, but rather that VWM passes single, attended items into LTM for comparison to the list of targets (Drew et al., 2016). Furthermore, in VWM studies, the CDA plateaus at VWM load of 3-4 items (Luria et al., 2016; Vogel & Machizawa, 2004). Our results may be interpreted to reflect that not all items of the MMS need to be loaded in VWM during memory search. Presumably, only “suspicious” targets from the memory set that are somewhat similar to the search item may fill up VWM (see Figure 1). The higher the MSS, the more likely is the search item to resemble more targets from the MSS. However, dissimilar targets will not be loaded into VWM; thus, the physical MSS would not be the number of targets actually loaded into VWM. This may also explain why the CDA was smaller and the MSS effect was absent in target absent trials. In the large set of distinct objects, there was overall small similarity between targets and distractors. Thus, most non-targets may have been rejected already on an attentional stage, not eliciting a memory search.

Indeed, the N2pc was only distinct for target present trials, suggesting attentional processing of targets but little attentional processing of non-targets. The present results are in agreement with recent hybrid search experiments (Lavelle et al., 2023), showing that feature-based attention can persist for target numbers exceeding VWM capacity. The N2pc was not significantly modulated by MSS. One might have expected that the N2pc would decrease with MSS, given that attentional guidance is assumed to be more effective when the number of search templates smaller (Ort & Olivers, 2020). Others have shown that the N2pc is smaller when observers search for two targets relative to only one target (Grubert et al., 2016; Grubert & Eimer, 2016). Note, however, that the design of the present task minimized the role of guidance by pre-cueing the target location from the outset. This N2pc does not reflect guidance, but only attentional processing of an item at an already selected location.

## Experiment 2

The findings of Experiment 1 suggest that the involvement of both VWM and LTM increases with target load up to eight target items in memory search. In Experiment 2, we sought to replicate and extend our findings with MSSs up to 64. Observers searched for 2, 4, 16, or 64 targets. We analyzed response times and accuracy, together with the N2pc, CDA, FN400, LPC and old/new effects, as a function of MSS and target presence. First, we expected to replicate that the amplitude of the FN400 amplitude and latency of the old/new effect increase with target load, marking prolonged recognition of targets with growing memory sets. Second, we tested whether, as in Experiment 1, the CDA amplitude would increase as MSS exceeded 4, here increasing to 16 and 64 targets. This would suggest an increase in VWM usage with target load, however, not following the item-specific capacity limitations in VWM. Finally, we expected that an N2pc should be elicited in target present trials as in Experiment 1, marking spatial attentional processing of the target item.

### Method

#### Participants

Thirty-nine observers from the University of Utah participated in the study in exchange for course credit or $15.00 an hour. Like in Experiment 1, we aimed for a sample size of 20 after exclusion, but overshot this goal. Ten participants had artifact rejection-rates that exceeded our exclusion criteria (30%) and one participant did not complete the study, leaving 28 participants in the final dataset (10 female/1 non-binary, average age: 22.8 years, age range: 18-36 years). The study was approved by the University of Utah’s Institutional Review Board, and all participants provided informed consent.

#### Memory Search Task

In Experiment 2, the MSSs were 2, 4, 16, and 64. Participants were required to pass the testing phase twice with a score of 80%. There were 296 trials for each MSS (50% target present, 50% target absent) in the memory search phase, and the task-relevant color was counterbalanced between experiment blocks rather than participants. Otherwise, the experimental design was identical to Experiment 1.

#### EEG Procedure and Analysis

The EEG procedure and analyses were identical to Experiment 1. On average, 14.9% of trials were rejected in the stimulus-locked waveforms and 12.6% of trials were rejected in the response-locked waveforms.

### Results

#### Response Time and Accuracy

Response time increased roughly logarithmically with MSS, F(3, 81)=15.54, p<.001, η_p_^2^=.37, BF_10_=7.01e+10 (Figure 8a). Response time differed between all set sizes, p<.0001, except between set size 2 vs. 4, p>.71. Response time for target present trials was significantly faster than target absent trials, F(1, 27)=33.85, p<.001, η_p_^2^=.55, BF_10_=2582.51. The set size by target presence interaction was not significant, F(3,81)=0.49, p=.69, η_p_^2^=.02, BF_10_=.06.

**Figure 8.**
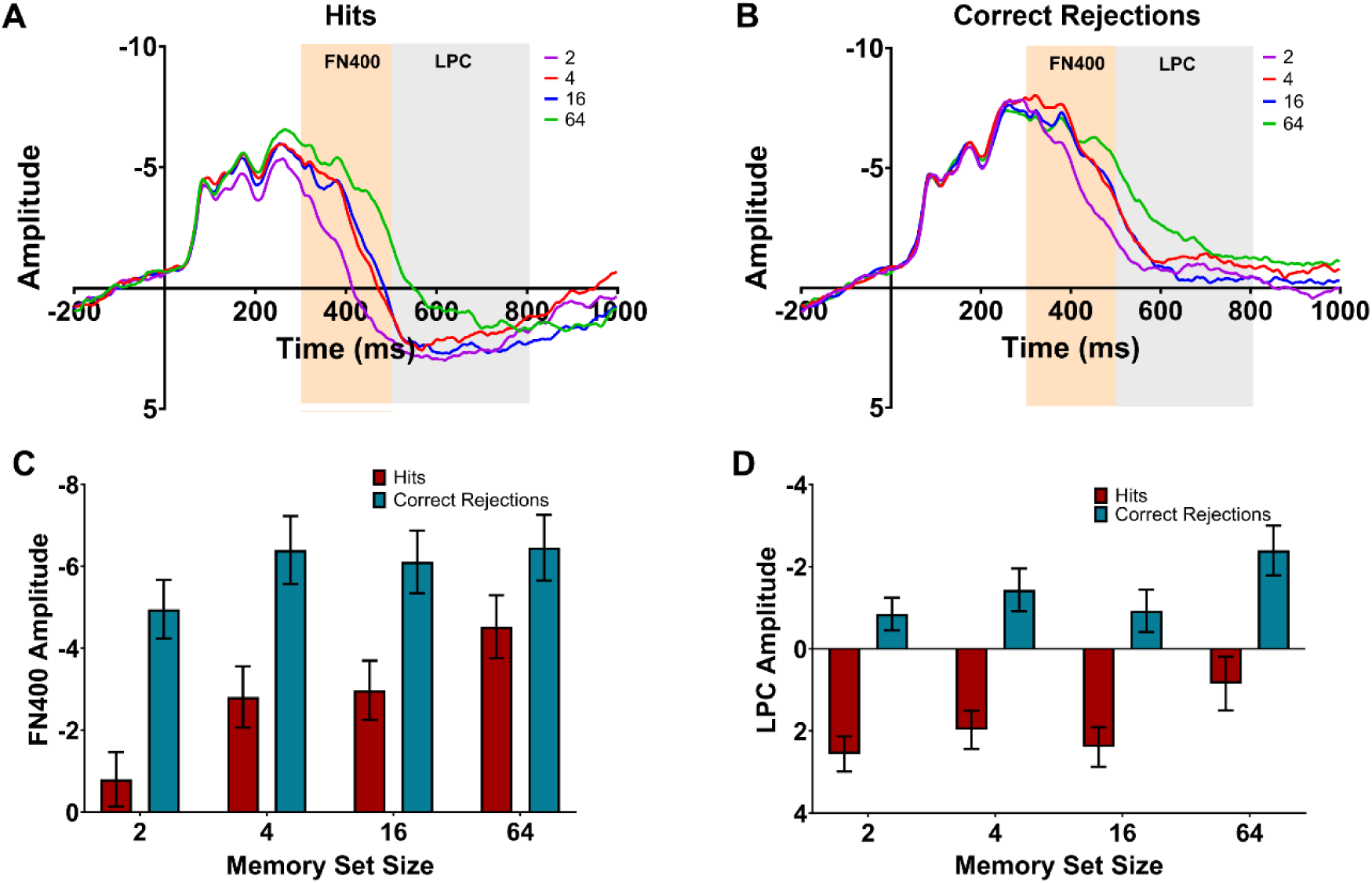
Non-lateralized data for Experiment 2: A) non-lateralized waveforms for target present trials. B) non-lateralized waveforms for target absent trials. C) mean FN400 amplitude. D) mean LPC amplitude.

Accuracy decreased with MSS, F(3, 81)=30.66, p<.001, η_p_^2^=.53, BF_10_=3.06e+16 (Figure 8b). Observers made significantly more errors when set size 64 was compared to the other set sizes, p<.0001. Accuracy was significantly lower for target present trials than target absent trials, F(1, 27)=7.5, p=.01, η_p_^2^=.22, BF_10_=2.59. The MSS by target presence interaction was not significant, F(3,81)=2.04, p=.11, η_p_^2^=.07, BF_10_=.17.

#### FN400, LPC and old/new effect

First, we evaluated the non-lateralized ERPs (Figure 8). As expected, we replicated the old/new difference, that is, more positive amplitudes for “old” targets compared to “new” distractor items, in the FN400, (1, 27)=209.4, p<.001, η_p_^2^=.89, BF_10_=9.43e+22, and the LPC, F(1, 27)=138.5, p<.001, η_p_^2^=.83, BF_10_=4.92e+27.

Furthermore, we replicated the FN400 modulation by MSS, F(3, 81)=15.31, p<.001, BF_10_=30888.11 (Figure 8c). The FN400 was significantly smaller for MSS 2 than MSSs 4, 16, and 64, all p-values<.001. There were no significant differences between any of the other MSSs, all p-values>.05, after correcting for multiple comparisons. The set size by target presence interaction was statistically significant, F(3,81)=15.72, p<.001, η_p_^2^=.37, BF_10_=10.97. In target present trials, the FN400 increased with increasing set size; all comparisons were significant, p<.0001, except the difference between set size 4 and 16. For target absent trials, the FN400 amplitude at set size 2 was significantly smaller than for set size 4, 16, and 64, p <. 0001, but the other set sizes did not differ significantly, p>.62.

The LPC amplitude also varied with MSS, F(3, 81)=6.99, p<.001, η_p_^2^=.21, BF_10_=12.49 (Figure 8d). Specifically, the LPC was significantly smaller (i.e. less positive) for the largest MSS of 64 compared to MSSs 2, 4, and 16, all p-values<.05. There were no significant differences between any of the other MSSs, all p-values>.05. The MSS by target presence interaction was not statistically significant, F(3, 81)=0.13, p=.94, η_p_^2^=.004, BF_10_=.06.

Finally, we also found that the mean amplitude of the old-new difference waveforms varied with MSS, F(3, 81)=4.37, p=.007, η_p_^2^=.14, BF_10_=5.70 (Figure 9b). MSS 64 had a significantly smaller amplitude than MSS 2, p=.03. None of the other MSS comparisons were statistically significant, all p-values>.05. Replicating the finding of Experiment 1, MSS modulated the fractional area latency of the old-new difference waves, F(3, 81)=12.11, p<.001, η_p_^2^=.31, BF_10_=16566.40 (Figure 9c). Fractional area latency was significantly shorter for MSS 2 than 16, p=.03, and 64, p<.001, shorter for MSS 4 than 64, p=.02, and shorter for MSS 16 than 64, p<.001.

**Figure 9.**
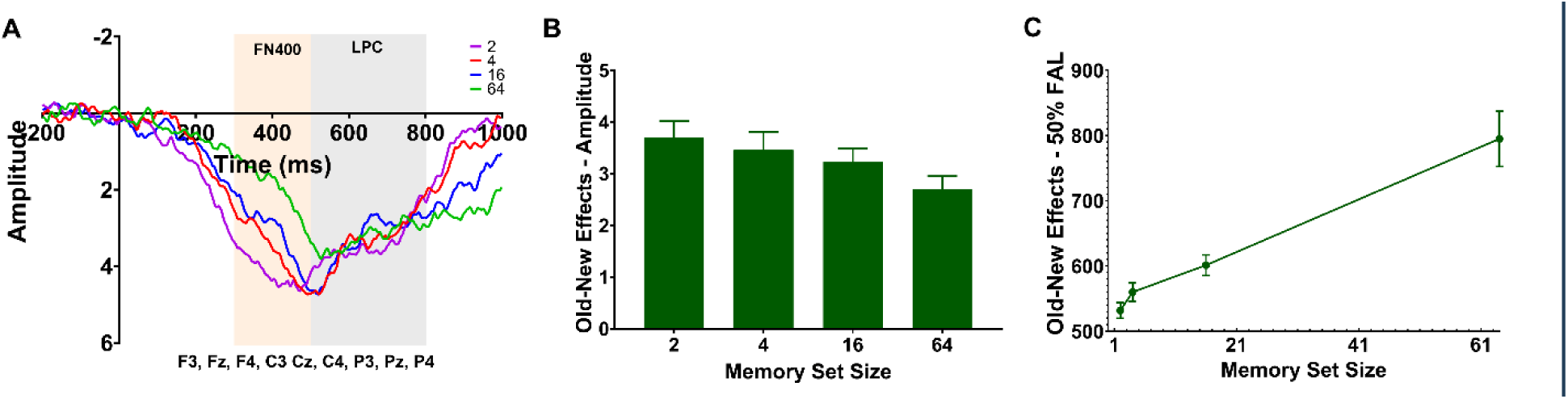
Old/new difference waves for Experiment 2: A) old/new difference waveforms B) mean amplitude of the old/new difference wave C) mean fractional area latency of the old/new difference wave

#### CDA

Next, we evaluated the stimulus-locked CDA (Figure 10). In line with our findings in Experiment 1, the grand-averaged ERPs suggest that the CDA continuously increased with MSSs. Indeed, the CDA was significantly modulated by MSS, F(3, 81)=4.18, p=.008, η_p_^2^=.13, BF_10_=.86 (Figure 10d), but post-hoc tests showed that the CDA was only significantly smaller (i.e., less negative) for MSS 2 than for MSS 64, p=.01. There were no significant differences between any of the other MSSs, all p-values>.05, after correcting for multiple comparisons. The CDA amplitude was larger for target present than target absent trials, F(1, 27)=10.77, p=.003, η_p_^2^=.29, BF_10_=42.54, and, different from Experiment 1, the MSS by target presence interaction was not statistically significant, F(3,81)=0.27, p=.85, η_p_^2^=.01, BF_10_=.07; thus, the effect of target load was comparable for target present and target absent trials.

**Figure 10.**
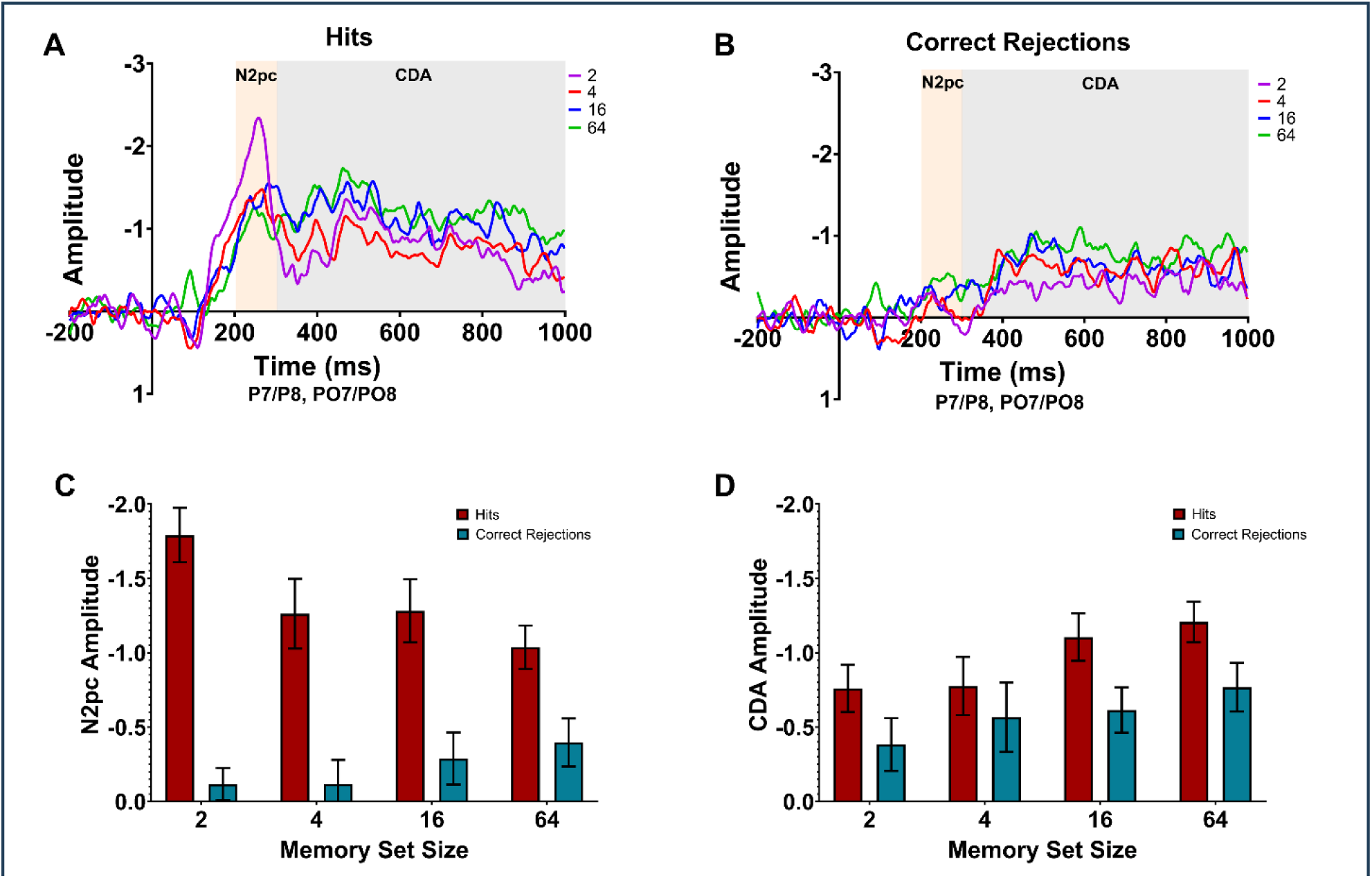
Lateralized, stimulus-locked data for Experiment 2: A) contralateral-ipsilateral waveforms for target present trials. B) contralateral-ipsilateral waveforms for target absent trials. C) mean N2pc amplitude. D) mean CDA amplitude.

#### N2pc

As in Experiment 1, the N2pc was prominent in target present, but not in target absent trials, F(1, 27)=104.6, p<.001, η_p_^2^=.79, BF_10_=3.31e+17. Again, the main effect of MSS on the N2pc amplitude was not significant, F(3, 81)=1.11, p=.35, η_p_^2^=.04, BF_10_=.06 (Figure 10c). However, the MSS by target presence interaction was statistically significant, F(3,81)=5.23, p=.002, η_p_^2^=.16, BF_10_=6.55. For target present trials, the N2pc was larger (i.e., more negative) for MSS 2 than MSSs 4, 16, and 64, all p-values<.05. No differences between MSSs were significant for target absent trials, all p-values>.05.

### Discussion

In Experiment 2, we largely replicated findings from Experiment 1 using larger MSSs. The FN400 and the latency of the old/new effect gradually increased with MSS. In addition, we found the LPC and old/new effect to be reduced for the largest set size of 64. These findings suggest that the recognition signal decreases, and occurs later, with increasing target load during memory search. Presumably, familiarity-based recognition is weaker because larger perceptual and semantic overlap between targets and non-targets in large target sets of distinct, highly variable, objects cause interference. This may also lead to a higher decision threshold for making old/new judgments, as more perceptual evidence needs to be accumulated to classify an item as target or non-target. Furthermore, higher error rates in the conditions with 64 targets suggest that participants may also have guessed correctly on a number of trials without a reliable recognition signal being present.

Interestingly, also the CDA increased, again; here with target load up to MSS 64, well beyond the item-capacity limits of VWM. This supports our assumption that the involvement of VWM during memory search cannot be understood simply in terms of a limited number of “spots” being filled with any target from LTM during the process of matching a visually selected object to targets from the memory set. Different from Experiment 1, the CDA also increased with MSS in target absent trials. As mentioned above, due to the perceptual and semantic overlap between targets and non-targets we can expect at larger MSS, “suspicious” non-targets might be matched against more targets from the MSS that become loaded into VWM.

As before, the N2pc was only observable in target present trials, even for larger MSSs, supporting again that some selective attention to relevant features even exists with large numbers of targets (Lavelle, Luria, & Drew, 2023). Furthermore, we found a larger N2pc for MSS 2 compared to larger MSSs. This might reflect that, at set size 2, observers could pre-activate two, relatively distinct, search templates, which facilitates attentional processing of the target if the MSSs is within the limits of VWM capacity (Ort & Olivers, 2020). At higher set sizes, perhaps only one or a few templates were preactivated.

## Experiment 3

Having established electrophysiological correlates of memory search in Experiments 1 and 2, we sought to use these measures to test predictions from Cunningham & Wolfe’s (2014) model of hybrid search in Experiment 3. A key assumption of that model is that memory search is not needed if items can be ruled out by attention to their categorical features (Shang et al., 2024). For example, when searching for multiple cats simultaneously, attention will be guided towards cats and items with similar features (e.g., dogs). These ‘lure’ items will elicit a memory search and will only be rejected during the target verification stage. In contrast, items that share few visual features in common with the cat target items (e.g., tables) will be rejected at an earlier stage and will not require a memory search (see Figure 1).

The CDA results in Experiment 1 and 2 suggest that VWM may indeed be filled selectively, only with “suspicious” targets from the MSS that are somewhat similar to the search item. In Experiment 3, we manipulated the MSS and the similarity between the search item and target category in order to test this assumption directly. Observers searched for 2 or 16 target objects from one category (e.g. cats), either among distractors of the same (other cats), a similar (dogs), or dissimilar (tables) category. If target-dissimilar items can be rejected before eliciting a memory search (Cunningham & Wolfe, 2014), we would expect that MSS influences the ERP correlates of memory search only if the attended item is of the same or a similar category as the target items. Accordingly, we expected that the FN400 and CDA would increase with MSS for targets and non-targets with similar categorical features, while no FN400 and CDA modulation would be found if observers attend to non-targets from a dissimilar category. Furthermore, we expected that targets and target-similar non-targets would elicit a larger N2pc than target-dissimilar non-targets, marking attentional processing of items with categorical target features (Shang et al., 2024).

### Method

#### Participants

Twenty-five participants from the University of Utah participated in the study for course credit or $15.00 an hour. Three participants exceeded the artifact rejection-rate threshold (30%) and two participants experienced equipment failure, leaving a total of 20 participants in the final dataset (13 female, 7 male, average age: 20.1 years, age range: 18-34 years), matching the pre-registered sample size (https://osf.io/tmurs)

#### Memory Search Task

The task structure of Experiment 3 was the same as Experiments 1 and 2 (Figure 11). However, in Experiment 3, participants memorized target sets containing objects from the same category. The MSSs were 2 and 16, and the possible target categories were dogs, cats, tables, and dressers. Each participant had one target set from the animal categories (e.g., dogs) and one target set from the furniture categories (e.g., tables) in separate blocks. Target categories and MSSs were counterbalanced across participants, such that there were an equal number of each category per set size. In the old/new recognition memory test, the foil items were from the same category as the target set for that block. During the memory search phase, the attended item varied between four conditions: a target (e.g., a target dog), a non-target from the target category (e.g., a non-target dog), a non-target from a similar category (e.g., a non-target cat), and a non-target from a dissimilar category (e.g., a non-target dresser). Each participant completed 174 trials per condition.

**Figure 11.**
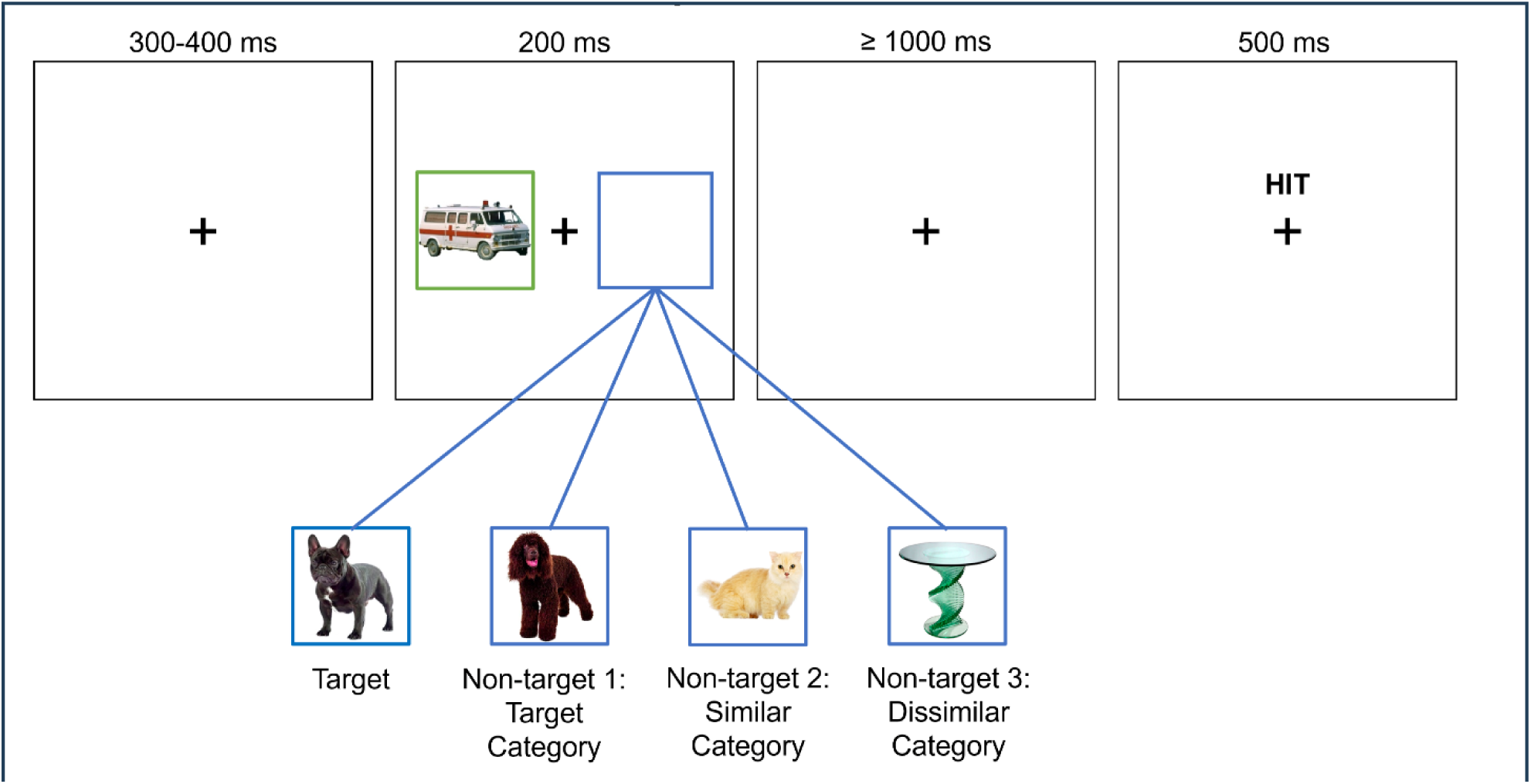
Experimental design for Experiment 3.

#### EEG Procedure and Analysis

The EEG procedure and analyses were identical to Experiments 1 and 2. 12.9% of trials were excluded from the stimulus-locked analysis and 9.6% of trials were excluded from the response-locked analysis.

#### Statistical Analysis

The analyses were pre-registered: (https://osf.io/tmurs). After collecting data from the first few participants, it became apparent that a categorical hybrid search task at MSS 64 would result in insufficient correct trials for our EEG analyses. Thus, we modified our pre-registration to include smaller MSSs and more trials. We performed a 2 (set size 2, set size 16) x 4 (Conditions 1-4) repeated measures ANOVA on each of the dependent variables. Main effects and interactions were followed-up with Tukey’s multiple comparisons.

### Results

#### Response Time and Accuracy

Response time decreased as the search item became more dissimilar to the target, F(3, 57)=52.00, p<.001, η_p_^2^=.73, BF_10_=234501.56 (Figure 12a). Specifically, response times were longer for targets than non-targets from the similar category, p<.001, and non-targets from the dissimilar category, p<.001. Response times were slower for non-targets from the target category than non-targets from the similar category, p<.001, and non-targets from the dissimilar category, p<.001. Response times were overall faster for MSS 2 than MSS 16, F(1, 19)=16.87, p=.001, η_p_^2^=.47, BF_10_=3.24e+8, and the search item condition by MSS interaction was significant, F(3, 57)=31.69, p<.001, η_p_^2^=.63, BF_10_=40.02. The MSS effect was significant for the target condition, p<.001, and the target category condition, p<.001, and marginally significant for the similar category, p=.06. The MSS effect was not significant for the dissimilar category, p=.43.

**Figure 12.**
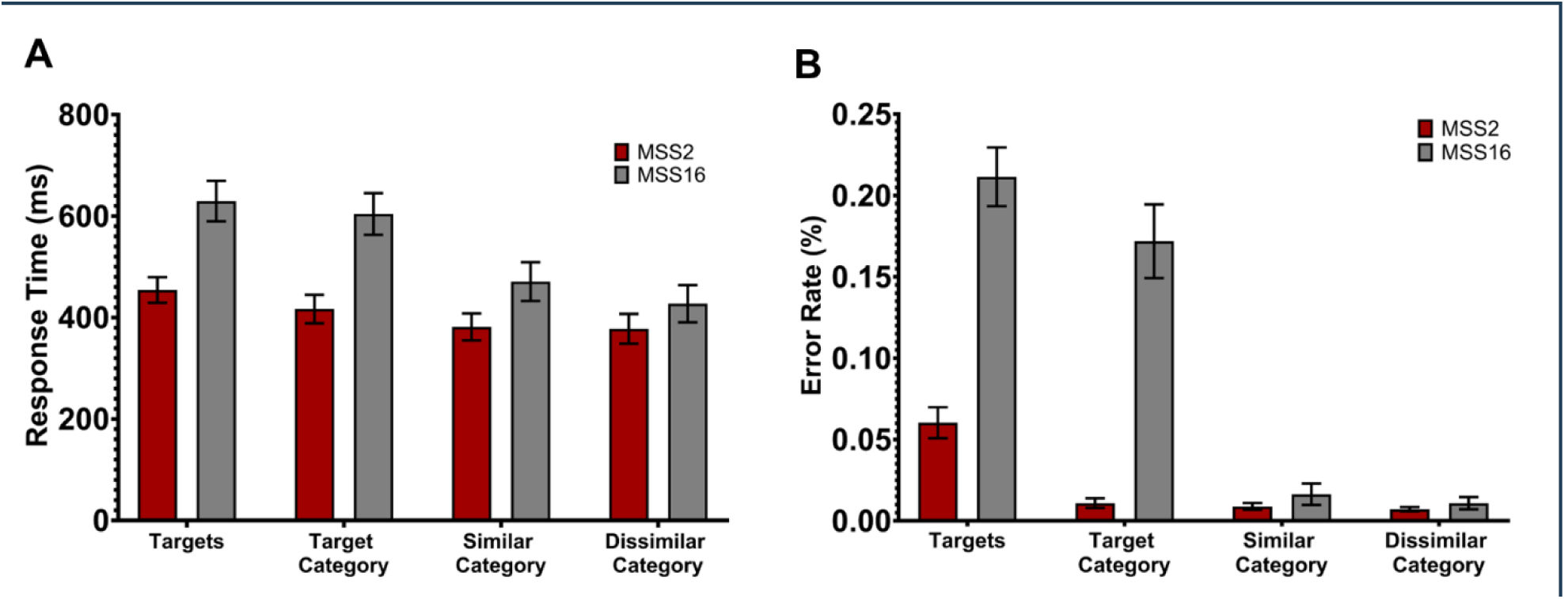
Behavioral data for Experiment 3: A) response time. B) error rate.

Accuracy increased as the search item became more dissimilar to the target, F(3, 57)=75.90, p<.001, η_p_^2^=.80, BF_10_=2.14e+12 (Figure 12b). Specifically, participants made more errors in classifying targets compared to non-targets from the target category, p<.001, non-targets from the similar category, p<.001, and non-targets from the dissimilar category, p<.001. Error rates were also higher for non-targets from the target category than for non-targets from the similar category, p<.001, and for non-targets from the dissimilar category, p<.001. Accuracy did not significantly differ between non-targets from the similar and the dissimilar category, p=.99. Accuracy was overall significantly better for MSS 2 than MSS 16, F(1, 19)=65.12, p<.001, η_p_^2^=.77, BF_10_=4.96e+6, and the search item condition by MSS interaction was significant, F(3, 57)=53.69, p<.001, η_p_^2^=.73, BF_10_=6.03e+14, such that the effects of categorical similarity were more pronounced at the higher MSS. MSS effects were significant for the target condition, p<.001, and the condition with distractors from the target category, p<.001. MSS effects were not significant for the condition with distractors from a target-similar category, p=.7, or the target-dissimilar category, p=.82.

#### FN400, LPC and old/new effects

First, we evaluated the non-lateralized waveforms (Figure 13). As in Experiment 1 and 2, the amplitude of the FN400 and LPC differed between “old” target items and “new” non-target items, F(3, 57)=38.71, p<.001, η_p_^2^=.67, BF_10_=9.92e+14 and F(3, 57)=55.18, p<.001, η_p_^2^=.74, BF_10_=2.29e+16 (Figure 13c, 13d). The FN400 and LPC were more positive for targets compared to all non-target conditions p<.0001; while the FN400 and LPC between non-targets of the target category, similar category, and dissimilar category did not differ significantly from each other after adjusting for multiple comparisons, all p-values>.08.

**Figure 13.**
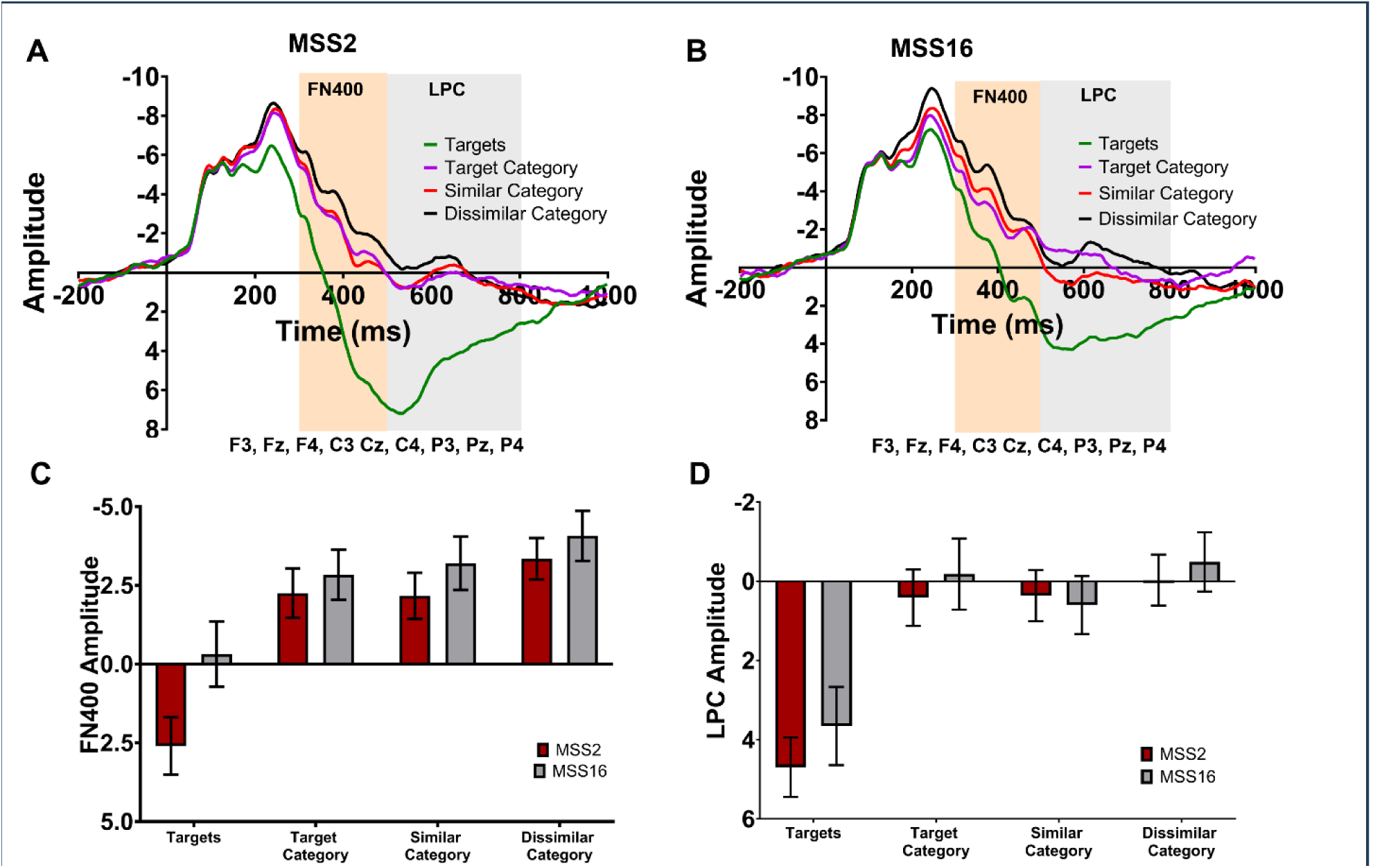
Non-lateralized data for Experiment 3: A) non-lateralized waveforms for memory set size (MSS) 2. B) non-lateralized waveforms for MSS 16. C) mean FN400 amplitude. D) mean LPC amplitude.

According to Cunningham & Wolfe (2014), we assumed that memory search is only elicited if the attended item shared categorical features with the target category; thus we would expect that the FN400 would vary with MSS if the search item is a member of the target or target-similar category, but not if the search item is a member of a target-dissimilar category. Indeed, the FN400 varied with MSS, F(1, 19)=8.35, p=.009, η_p_^2^=.31, BF_10_=8.32 (Figure 15c) and MSS interacted significantly with search item condition, F(3, 57)=5.76, p=.002, η_p_^2^=.23, BF_10_=1.40. However, the FN400 was only significantly larger (i.e. more negative) for MSS 16 than MSS 2 in the target condition, p=.001, while the MSS effect was not significant for any of the non-target conditions, all p-values>.05. In accordance with the findings from Experiment 1, the LPC amplitude did not significantly vary with MSS, F(1, 19)=0.91, p=.35, η_p_^2^=.05, BF_10_=.30. and neither was the condition by MSS interaction significant, F(3, 57)=0.93, p=.41, η_p_^2^=.05, BF_10_=.13 (Figure 13d).

The amplitude of the old-new recognition effects (Figure 14) decreased from MSS 2 to 16, F(1, 19)=4.79, p=.01, η_p_^2^=.20, BF_10_=136.16, and differed significantly between non-target conditions, F(2, 38)=6.07, p=.02, η_p_^2^=.24, BF_10_=.47. Specifically, the old-new difference was larger for target-dissimilar non-targets than for target-similar non-targets, p=.02, and marginally significantly larger for target-dissimilar non-targets than for non-targets of the target category, p=.06. The condition by MSS interaction was not significant, F(2, 38)=0.14, p=.87, η_p_^2^=.01, BF_10_=.14. Also the fractional area latency of the old-new effects significantly differed between non-target conditions, F(1, 19)=9.07, p=.007, η_p_^2^=.32, BF_10_=.12 (Figure 16d), but none of the post-hoc tests survived the correction for multiple comparisons, all p-values>10. The main effect of MSS on the fractional area latency was not significant, F(2, 38)=0.80, p=.46, η_p_^2^=.04, BF_10_=1402.10; however, MSS interacted with the non-target condition, F(2, 38)=3.44, p=.04, η_p_^2^=.15, BF_10_=.51. Specifically, the old-new effect occurred later for MSS 16 than 2 in the difference wave between targets and non-targets of the target category, p<.001, and in the difference wave between targets and non-targets from a dissimilar category, p<.001.

**Figure 14.**
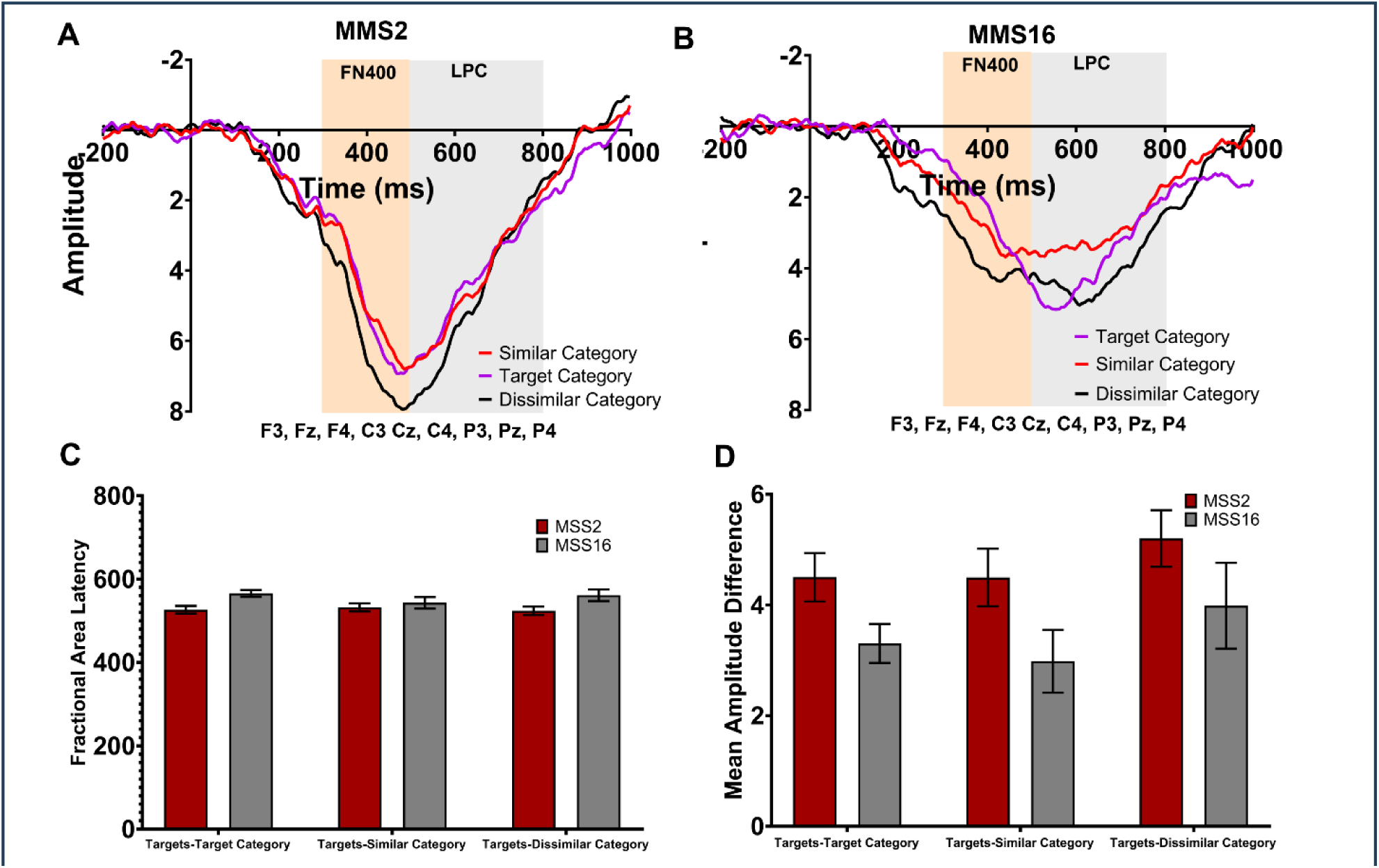
Old/new difference waves for Experiment 3: A) old/new difference waveforms for memory set size (MSS) 2 B) old/new difference waveforms for MSS 16 C) mean amplitude of the old/new difference wave C) mean fractional area latency of the old/new difference wave.

#### CDA

Next, we evaluated the stimulus-locked CDA (Figure 17). As in Experiments 1 and 2, the CDA amplitude increased significantly with MSS, F(1, 19)=19.92, p<.001, η_p_^2^=.51, BF_10_=69.90, confirming again that the involvement of VWM increases with target load during memory search. If VWM involvement during this memory search is indeed restricted to items that share categorical features with the target (Cunningham & Wolfe, 2014), we would expect that the CDA would be more pronounced and would vary with MSS only for targets and non-targets from the same or similar category. Indeed, the CDA varied with similarity between the attended item and the target category, F(3, 57)=21.52, p<.001, η_p_^2^=.53, BF_10_=8.93e5 (Figure 15d), and increased gradually with similarity to the target: CDA amplitudes were larger for targets than non-targets from the similar category, p<.001, and the dissimilar category, p<.001, and larger for non-targets from the target category than non-targets from the similar category, p=.004, and non-targets from the dissimilar category, p<.001. The CDA amplitude did not significantly differ between non-targets from the similar and dissimilar category, p=.30. Furthermore, the search item condition by MSS interaction was significant, F(3, 57)=7.23, p=.004, η_p_^2^=.28, BF_10_=413.39. The difference between MSSs 2 and 16 was significant for targets, p<.0001, and items from the target category, p=.0005, but not for items from a similar, p=.14, and dissimilar category, p=.18.

**Figure 15.**
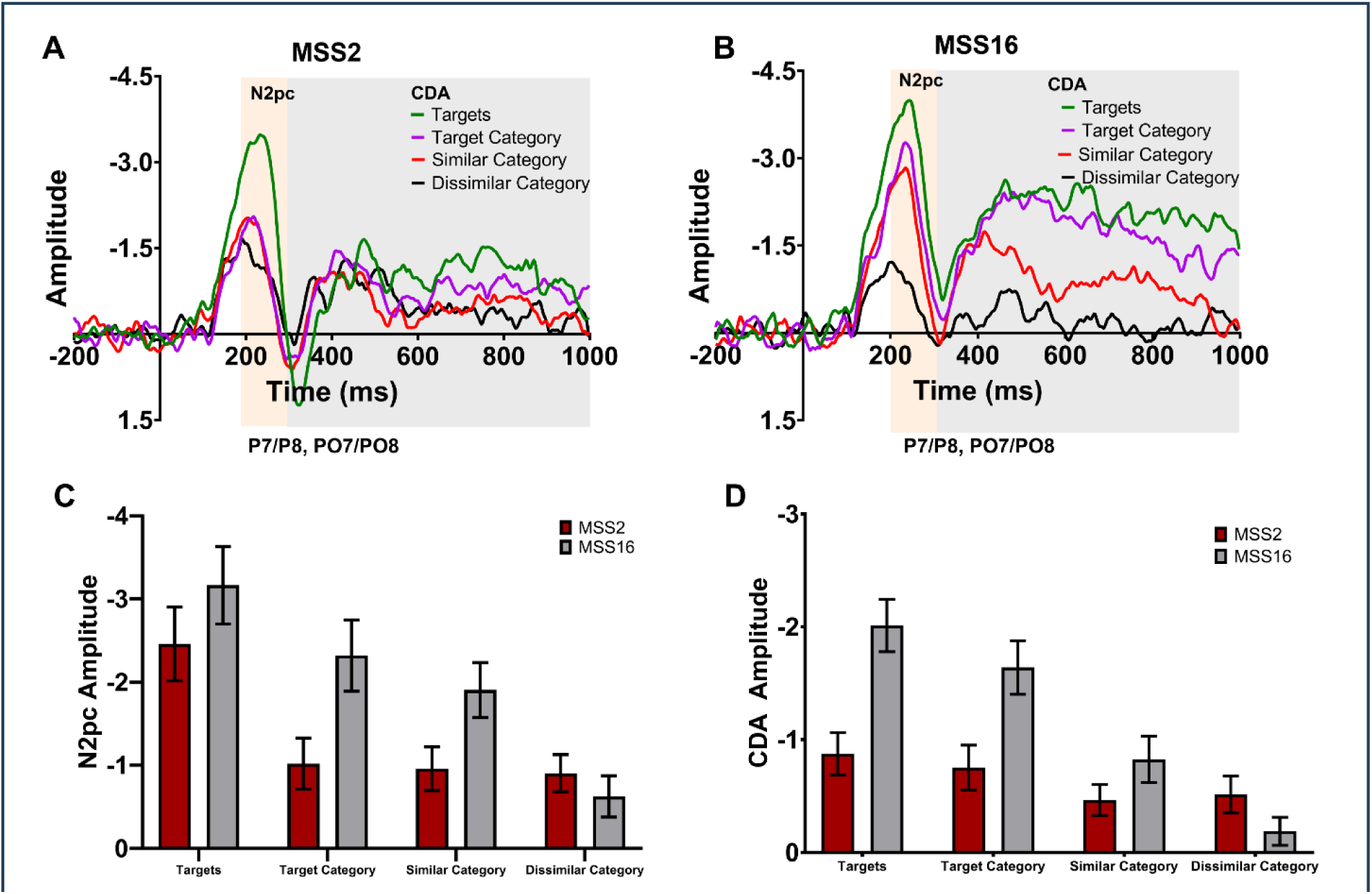
Lateralized data for Experiment 3: A) contralateral-ipsilateral waveforms for memory set size (MSS) 2. B) contralateral-ipsilateral waveforms for MSS 16. C) mean N2pc amplitude. D) mean CDA amplitude.

#### N2pc

Finally, we expected that attentional processing of categorical target features would be marked by the N2pc (Shang et al., 2024). Indeed, the amplitude of the N2pc (Figure 15) significantly decreased (became less negative) as the search item became more dissimilar to the target, F(3, 57)=16.20, p<.001, η_p_^2^=.46, BF_10_=3.37e+8 (Figure 17c). Specifically, the N2pc was larger for targets relative to non-targets from the target category, p=.002, non-targets from the similar category, p<.001, and non-targets from the dissimilar category, p<.001. The N2pc was also larger for non-targets from the target category than non-targets from the dissimilar category, p=.02. Furthermore, the N2pc was significantly larger for MSS 16 than MSS 2, F(1, 19)=17.16, p=.001, η_p_^2^=.47, BF_10_=13.02, and the condition by MSS interaction was significant, F(3, 57)=7.28, p=.002, η_p_^2^=.28, BF_10_=4.16. The MSS effect was significant for the targets, p=.02, non-targets from the target category, p<.001, and non-targets from the similar condition, p=.002. MSS effects were not significant for non-targets from the dissimilar category, p=.72.

### Discussion

In Experiment 3, we replicated that the FN400 and the CDA are sensitive to target load during memory search. In addition, our results support the assumption that target-similar items elicit a memory search, while target-dissimilar items can be rejected prior to engaging in memory search (Cunningham & Wolfe, 2014; Lavelle, Luria, & Drew, 2023; Shang et al., 2024): Observers responded more slowly and less accurately to search items that are more similar to the target; particularly when the MSS was larger. These behavioural costs of target similarity and MSS were mirrored in multiple ERP variations: First, the CDA increased with MSS if the search item was a target or a non-target from the same target category, which indicates that items with target-matching categorical features activate VWM resources during memory search, while categorically different items put less (or no) load on VWM. Furthermore, the FN400 increased with MSS if the search item was a target and the old/new effect gradually decreased as the target-similarity of the non-target and MSS increased. Presumably, the LTM-based recognition is more difficult under high target-distractor similarity and target load due to perceptual and conceptual interference (Konkle et al., 2010). Distractors of the target category may elicit a certain degree of “lure” familiarity, specifically if the MSS is larger.

The N2pc was larger for targets and non-targets of the target category. The N2pc also increased with MSS for targets and non-targets of the target category, but not if the search item was from a target-dissimilar category. This finding is in line with, and extends, recent results (Shang et al., 2024) showing that attentional processing is enhanced for items that match features of the target category. This effect depends on the number of targets in memory: At smaller MSSs, observers could effectively prevent items from similar categories from being attentionally processed, just as they prevented dissimilar items from being processed. However, at larger MSSs, target-similar non-target items were more likely to be attentionally processed.

## General Discussion

In recent years, our understanding of search behavior has moved toward increasingly complex and realistic tasks (e.g., radiology) (Nartker, Alaoui-Soce, & Wolfe, 2020; Turoman et al., 2021; Wolfe, 2021). In our daily routines, we often perform “hybrid” visual and memory search; that is, the simultaneous search through the environment for multiple-items stored in memory (Wolfe, 2012). One real-world example of hybrid search is searching through your mental shopping list while looking for the items on your list at the grocery store (Boettcher, Drew, & Wolfe, 2018). Although the role of VWM and LTM processes has been studied in relatively simple visual search tasks (e.g., Woodman et al., 2007), it is unclear how different memory processes contribute to more complex hybrid search tasks, in which the MSS can be much higher with the possibility of hundreds of possible targets, held in memory. Here, we used EEG to better understand the mechanisms underlying the memory search component of hybrid search by manipulating target load (MSS). In Experiments 1 and 2, observers searched for sets of 1-64 distinct object images. We assumed that VWM load, as indexed by the CDA (Luria et al., 2016), would increase for memory search of small target sets within the capacity limitations of VWM. With MSSs significantly larger than 4, we expected to see a ceiling effect in measures of VWM. We expected that search through larger memory sets would rely on LTM recognition processes (Ort & Olivers, 2020), as indexed by the FN400, LPC, and old/new effect (Curran, 2000; Rugg & Curran, 2007). In accord with this second set of expectations, we found that the FN400 and old/new effect varied with MSSs up to 64 during memory search, presumably marking increased demands of LTM retrieval. However, our expectations about the CDA and VWM were not met. The CDA also increased with MSS up to 64, far beyond any reasonable estimate of the capacity of VWM. This suggests that VWM resources during memory search cannot be understood as a simple matter of filling up a small number of “slots”. In Experiment 3, we tested whether categorical similarity of targets and non-targets would influence VWM and LTM processes during memory search. We found electrophysiological support for the idea that memory search for targets derived from a single category can be limited to target-similar items (Cunningham & Wolfe, 2014; Shang et al., 2024). Items from target-dissimilar non-target categories can be dismissed without requiring a search through the memory set of specific targets. Target load effects on the FN400 and CDA depended on the categorical status of the search item. Furthermore, the N2pc increased with similarity to the target; in particular, when the MSS was large, suggesting that items dissimilar to the target category were rejected early, prior to memory search.

### The role of long-term memory in memory search

It has been proposed that in hybrid search tasks with large sets of well-learned target items, target verification is based on LTM (Drew et al., 2017; Ort & Olivers, 2020). Accordingly, the decrease in search performance with increasing MSS may reflect the “list length effect”, which refers to the phenomenon that longer lists of learned, to-be-recognized items are typically associated with poorer recognition memory performance compared to shorter lists (Shiffrin & Steyvers, 1997). The present experiments show that target load during memory search performance indeed modulated ERP markers associated with LTM recognition. Specifically, the FN400, assumed to mark familiarity-based LTM recognition (Rugg & Curran, 2007), was larger for “old” targets than “new” non-targets and increased gradually with MSS. Furthermore, the old/new difference was prolonged and decreased for large MSSs. Familiarity-based recognition is considered to be a fast and relatively automatic process (Curran, 2000), which may explain why search through many items in memory is remarkably quick (Wolfe, 2012). Consistent with this proposal are recent results showing that familiar items have an advantage over novel stimuli (Madrid et al., 2019) and increasing familiarity of non-targets and reducing familiarity of targets can cause some response time costs in hybrid search (Wiegand & Wolfe, 2020; Wolfe et al., 2015). The FN400 and old/new effect modulations suggest that the familiarity signal to verify an item as “old” becomes weaker and slower the more targets are in the memory set. Presumably, longer lists of distinct target objects cause more interference during the memory search due to larger conceptual and perceptual overlap between the target set and non-targets (Konkle et al., 2010). This is also supported by the findings of Experiment 3, in which the old/new difference between targets and target-similar non-targets is smaller than between targets and target-dissimilar non-targets. Categorical similarity is also associated with conceptual and perceptual overlap (Küper et al., 2012; Küper & Zimmer, 2018) that weakens the discriminative strength of the familiarity signal between targets and target-similar non-targets during memory search.

In contrast to familiarity-based recognition, recollection-based recognition is more time consuming, effortful, and less susceptible to interference by item similarity (Yonelinas, 2002). Behavioural studies have shown that hybrid search remains efficient even if targets cannot be identified based on a familiarity signal alone and a form of rapid recollection may support recognition (Guild et al., 2014; Wolfe et al., 2015). In the present study, we found that the LPC was little modulated by MSS, suggesting that recollection-based retrieval, while likely contributing to target recognition in hybrid search (Wolfe et al., 2015), remains largely undisturbed by the growing target load.

### The role of visual working memory and attentional processing in memory search

Our findings confirm that target load affects LTM recognition in hybrid search, but also provide compelling evidence that attentional and VWM processes contribute to memory search, even if the target memory set is (very) large. Our previous behavioral data suggested that VWM does not play a MSS-dependent role in hybrid search (Drew et al., 2016). Furthermore, memory search was proposed to be VWM-based for set sizes within VWM capacity limits, but LTM-based for set sizes beyond VWM capacity limits (Ort & Olivers, 2020). Accordingly, we hypothesized that the CDA, marking VWM usage (Luria et al., 2016), may plateau around MSSs of ∼3-4 items (Luck & Vogel, 1997). However, all experiments showed that the CDA continued to increase with MSS beyond the VWM capacity limit.

It is unlikely that the CDA increase reflects that many more than four target representations in VWM in the present task (Ikkai, McCollough, & Vogel, 2010; Vogel & Machizawa, 2004). Rather, the CDA modulation may reflect sustained effort while the search item is matched against multiple target templates retrieved from LTM. Presumably, only those targets that share features with the perceptually presented item will be loaded into VWM, but not all targets from the MSS. This explains that the CDA MSS effect does not strictly follow the physical set size of the target set and that the CDA is sensitive to the similarity of the non-targets. More generally, the CDA was shown to rise with difficulty of VWM operations in other tasks, such as mental rotation (Ankaoua & Luria, 2023). In the present tasks, one possibility is that a higher fidelity version of the attended item might be passed into VWM for comparison to the target memory set when target verification becomes more difficult, taking up more resources in VWM (Bays, Wu, & Husain, 2011). The high fidelity comparison process may also explain why distractors are better recognized when they resemble targets and MSS is larger (Lavelle et al., 2023).

Of note, as expected, the CDA (and FN400) modulations by MSS were observed both in target present and target absent trials. This is in line with the assumption that both targets and non-targets elicit a memory search in which target verification occurs on post-selective processing stages (Ort & Olivers, 2020). Notably, however, the CDA was larger and the MSS effect was more pronounced on target present trials than target absent trials. Possibly, in Experiments 1 and 2, where targets and non-targets were distinct object images, some (randomly) target-dissimilar non-targets might have been rejected easily, thus, were not loaded into VWM (Hilimire et al., 2011) and did not elicit a CDA. This would lead to lower CDA amplitudes on average in target-absent, as compared to target-present, trials. In Experiment 3, where target–non-target similarity was systematically varied, CDA amplitudes were indeed strongly reduced in response to target-dissimilar non-targets items, supporting that target-dissimilar items are not loaded into VWM.

Finally, the N2pc marks how attentional processing of the search items influences memory search. In line with previous studies, the N2pc amplitude decreased with increasing the number of targets (Grubert & Eimer, 2016), which was significant only in Experiment 2. Presumably, if the observer looks for one out of two targets, it is possible to activate two distinct search templates, with high fidelity, facilitating attentional processing of targets relative to non-targets. When looking for one target out of 8, 16, or 64, this search template will be crude so that attentional processing of targets and non-targets is more equal (i.e. less biased to specific features represented in a search template). Experiment 3 further demonstrated that the N2pc was sensitive to the degree of feature overlap between the search item and the target category (Shang et al., 2024). Targets and non-targets from the same category both elicited a strong N2pc, suggesting that attentional processing of target-similar non-targets contributes to the diminishment of recognition accuracy (Konkle et al., 2010) and hybrid search performance (Lavelle et al., 2023), and influences target verification on post-selection processes, as we see in the modulation of the CDA and FN400. In fact, the degree of interference due to list length (Criss & Shiffrin, 2004a) in memory search could also be understood as the ability to attentionally prioritize the memorized items, which is easier, or can be done with higher fidelity, if the number of items is smaller and feature overlap is lower.

Lastly, similar to the CDA, the N2pc was larger for target present than target absent trials. This lends support to recent findings showing that attentional prioritization is not absent even for larger number of unrelated targets (Lavelle et al., 2023). Of note, however, the experiments reported here were designed to examine the memory search component of hybrid search, rather than the attentional selection stage. Observers were only shown one item on the cued side of the lateralized display, greatly reducing the selective attention requirements in the first place. Thus, it remains unclear how these findings might interact with stronger demands on visual selection, such as increasing the visual set size, and this would be a compelling area for future research.

## Appendix: Analyses of the response-locked CDA

We analyzed the response-locked CDA (-300 to 0 ms), as opposed to the stimulus-locked CDA reported in the main text, to ensure any observed differences in mean amplitude between conditions were not driven by differences in response-time (Ankaoua & Luria, 2023; Williams & Drew, 2021).

### Experiment 1, memory set size (MSS) 1-8

The response-locked CDA in Experiment 1 is shown in Figure 1. Similar to the stimulus-locked CDA, the response-locked CDA amplitude varied with MSS, F(3, 57) = 2.81, p = .047, BF_10_ = .18, and was larger for target present than target absent trials, F(1, 19) = 21.31, p < .001, BF_10_ = 2.22e+6. However, the MSS by target presence interaction was not statistically significant, F(3, 57) = 1.2, p = .32, BF_10_ = .26, and none of post-hoc tests survived the adjustments for multiple comparisons, all p-values > .05.

**Figure 1.**
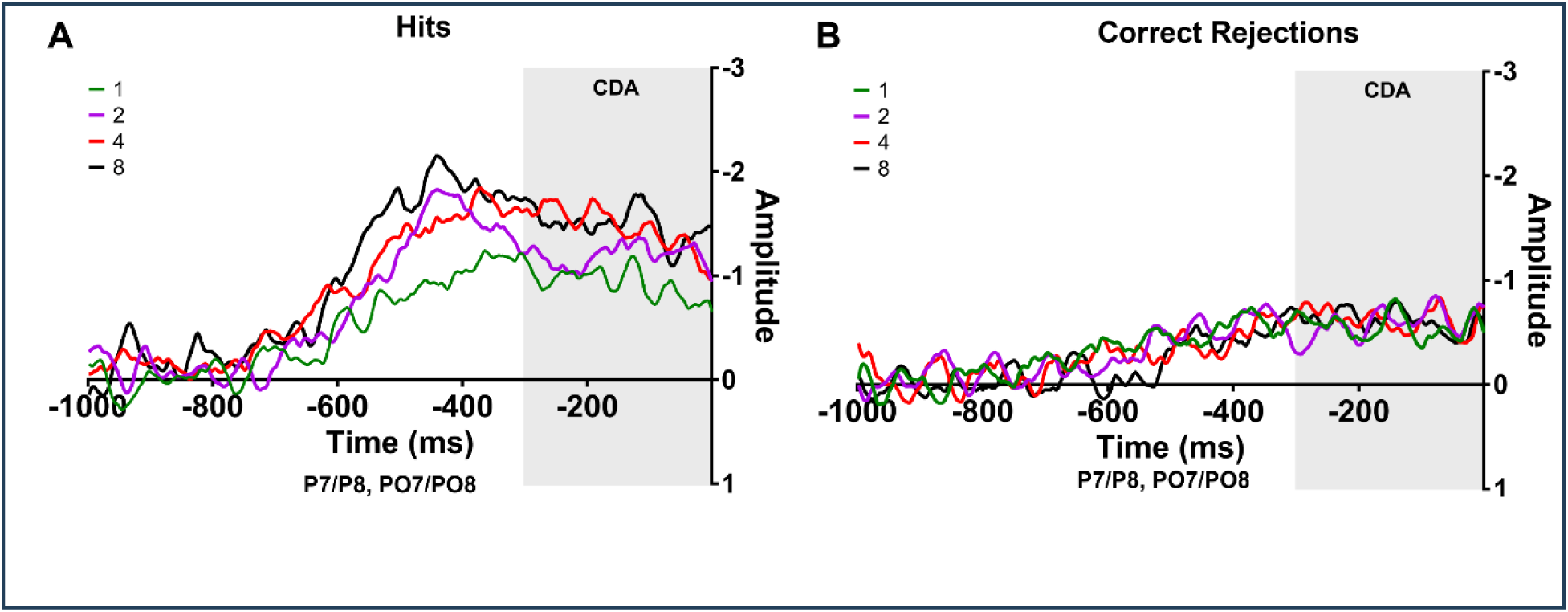
Response-locked lateralized waveforms for Experiment 1: A) target present trials, B) target absent trials.

### Experiment 2, MSS 2-64

The response-locked CDA in Experiment 2 is shown in Figure 2. Similar to the stimulus-locked CDA, the response-locked CDA amplitude also varied with MSS, F(3, 81) = 4.18, p = .008, BF_10_ = .70, so that the CDA amplitude was significantly smaller (less negative) for MSS 4 than for set size 64, p = .006. There were no significant differences between any of the other MSS comparisons, all p > .05. The main effect of target presence did not reach significance, F(1, 27) = 3.69, p = .07, BF_10_ = 1.23, and the set size by target presence interaction was not significant, F(3, 81) = .12, p = .95, BF_10_ = .06.

**Figure 2.**
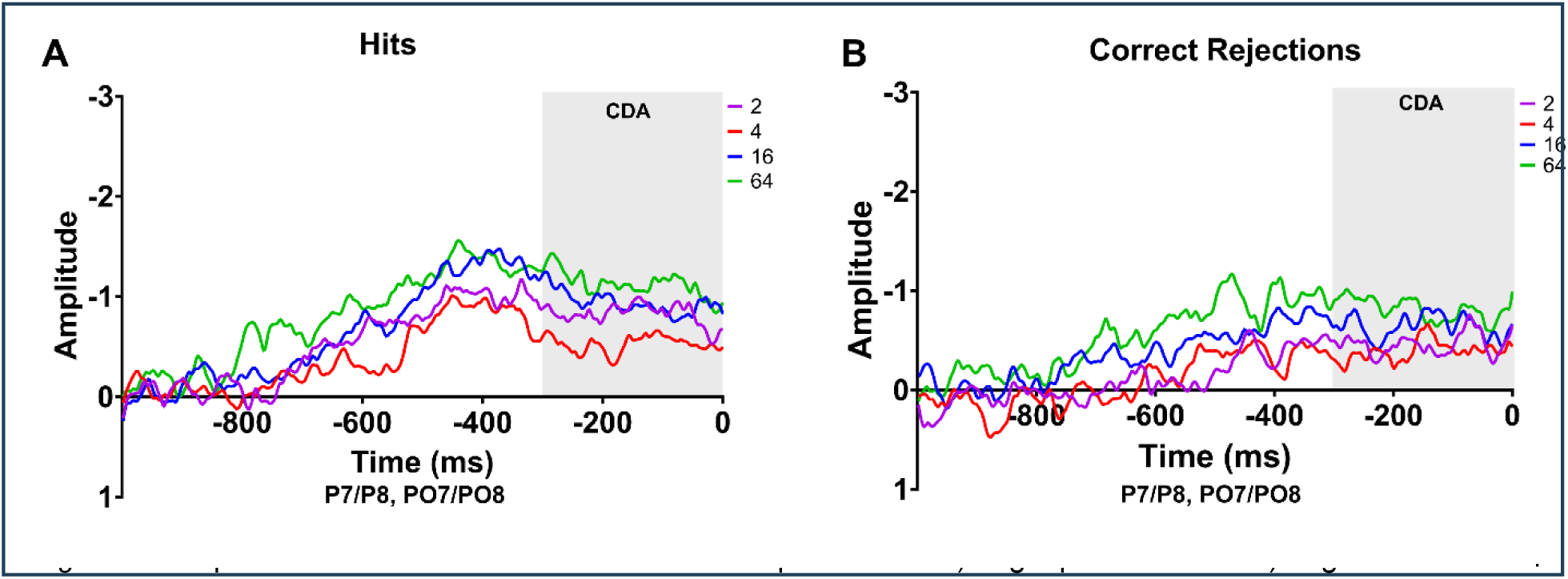
Response-locked lateralized waveforms for Experiment 2: A) target present trials. B) target absent trials.

### Experiment 3 categorical similarity, MSS 2 and 16

The response-locked CDA in Experiment 3 is shown in Figure 3. Similar to the stimulus-locked CDA, the amplitude of the response-locked CDA also increased with MSS, F(1, 19) = 16.86, p = .001, BF_10_ = 52.81, and varied with similarity between the attended item and the target category, F(3, 57) = 10.26, p < .001, BF_10_ = 4070.98. Similar to the stimulus-locked effects, the response-locked CDA was larger for the targets than non-targets from the similar category, p = .02, and non-targets from the dissimilar category, p < .001, and larger for non-targets from the target category than non-targets from the dissimilar category, p = .001. None of the other comparisons were significant, all p-values > .05. Also the condition by MSS interaction was significant, F(3, 57) = 7.15, p = .002, BF_10_ = 33.37, reflecting that the CDA increased with MSS for targets, p < .0001, and non-targets from the target category, p = .0002, but not for non-targets from a similar, p = .18, and dissimilar category, p = .30.

**Figure 3.**
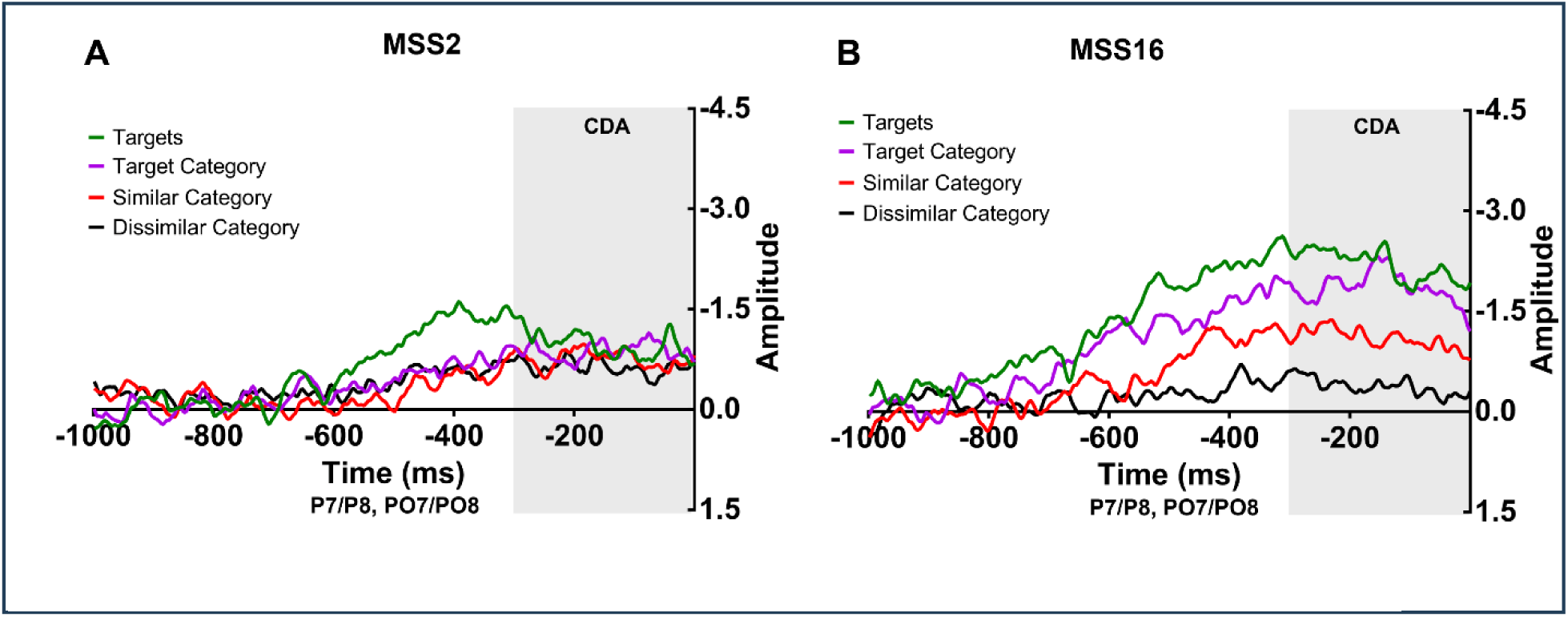
Response-locked waveforms for Experiment 3: A) memory set size (MSS) 2. B) MSS 16.

## Notes

**Funding:** This material is based upon work supported by the National Science Foundation Graduate Research Fellowship under Grant No. 1747505.

https://osf.io/cdp4f/

https://osf.io/dms4q/

https://osf.io/zcsxv/

### Competing Interest Statement

The authors have declared no competing interest.

